# TREM2 encodes plaque macrophage plasticity to mediate atherosclerosis regression

**DOI:** 10.64898/2026.10.31.763330

**Authors:** Hannah Hillman, Gage M. Stuttgen, Su-Jeong Lee, Michael T. Patterson, Michael D. Chang, Yingyue Zhou, Ainsley E. Kennedy, Patricia R. Schrank, Omer Mitha, Tessa J. Barrett, Marco Colonna, Jesse W. Williams

## Abstract

Cholesterol lowering therapies stabilize atherosclerotic plaque and reduce ischemic events, but the molecular mechanisms governing plaque remodeling or regression remain poorly defined. The lipid sensor TREM2 is a key modulator of plaque-associated foam cell development and was upregulated by plaque macrophages in atherosclerosis regression. We examined macrophage- or monocyte-specific *Trem2* deletion during regression in mice and found that TREM2 expression on plaque macrophages, not newly recruited monocytes, was required for plaque regression. *Trem2* deletion impeded necrotic core resolution, leading us to interrogate the efferocytic capacity of TREM2^+^ cells during lipid lowering. An *in vitro* model of cholesterol-experienced ex-lipid associated macrophages “Ex-LAMs” demonstrated that prior lipid priming enhanced efferocytic ability after unloading lipids and maintained expression of lysosomal and cytoskeletal remodeling genes. Augmented TREM2 signaling through genetic models or agonist treatment accelerated efferocytic ability in vitro and plaque regression in vivo. Thus, we posit that a TREM2-dependent ExLAM programming drives atherosclerosis resolution and facilitates beneficial plaque remodeling through enhanced efferocytosis. Overall, targeting the TREM2 pathway may be a viable strategy for stabilizing atherosclerotic plaques in patients treated with aggressive cholesterol control.

## INTRODUCTION

Atherosclerosis regression describes the reduction of plaque size and remodeling of plaque composition in response to the lowering of cholesterol^1^. In addition to reducing plaque burden, regression is associated with increased plaque stability, characterized by reduced necrotic core area, thicker fibrous cap, and overall reduced inflammation. Mechanisms mediating plaque regression remain contentious, with prior work suggesting monocyte entry is halted in regression, yet monocytes support the major regression phenotype^2,3^. With intense cholesterol lowering therapies, such as PCSK9 inhibitors, being increasingly used to treat hypercholesterolemic patients, it is now possible to achieve regression of human plaques^4^. However, plaque regression is not observed in all patients and the mechanisms governing the changes in plaque remodeling during regression remain incompletely understood.

Macrophages play a central role in both progression and regression of atherosclerotic plaque. During disease progression, cholesterol-loaded foam cells, a plaque-associated population of lipid-associated macrophage (LAM), emerge in response to lipid accumulation in the initma, contributing to the handling of excess cholesterol and the clearance of dead cells and debris, termed efferocytosis^5,6^. However, prolonged lipid accumulation in foam cells often leads to cellular dysfunction and apoptosis, overall driving the formation of necrotic cores. Similar observations have been associated with LAMs across other disease settings^7,8^. The LAM cellular program and dysfunction have been previously associated with a terminal differentiation state, in most cases leading to cell death and contributing to disease pathology. It is unknown whether promoting lipid clearance from foam cells or LAMs can mediate macrophage rejuvenation and whether these pathways could be targeted therapeutically to mediate beneficial plaque remodeling and promote atherosclerosis resolution.

LAMs have emerged as an attractive therapeutic target due to their role in driving tissue remodeling and immune suppression^9–11^. Prior studies have implicated Triggering Receptor Expressed on Myeloid Cells 2 (TREM2) as a potent regulator of local lipid load and cellular debris, modulating how LAMs respond to tissue damage in disease progression^7,12,13^. TREM2 supports cell survival while influencing lipid uptake, efflux, and efferocytosis, processes that shape plaque composition and stability^11,14^. Until now, the role of TREM2 in atherosclerosis regression has not been systematically investigated, and the question remains whether TREM2 plays a role in the adaptation of LAMs to a lower lipid environment.

In this study, we propose that plaque macrophages undergo a TREM2-mediated phenotypic switch in response to cholesterol lowering that drives atherosclerosis regression and plaque remodeling. Moreover, we define the formation of lipid experienced “ExLAMs” that possess enhanced efferocytic and proliferative capacity while maintaining expression of core LAM genes, including *Trem2*. Finally, we show that boosting TREM2 signaling through antibody agonism or cleavage resistance drives atherosclerosis regression and efferocytosis. In all, these findings demonstrate that targeting LAMs and ExLAMs through TREM2 signaling and lipid clearance may represent a viable therapeutic option for bolstering plaque stability in a clinical setting.

## RESULTS

### Myeloid-Specific Trem2 Deletion Impedes Atherosclerotic Plaque Regression and Necrotic Core Resolution

Given the known role of TREM2-expressing macrophages in promoting tissue remodeling and lipid handling, we sought to test the hypothesis that TREM2 is required for pro-resolving macrophage function during atherosclerosis regression. To model atherosclerosis regression, plaque-bearing *Ldlr*^-/-^ mice were administered a liver-targeted AAV to drive the expression of the murine *Ldlr* gene in hepatocytes, promoting rapid and sustained cholesterol control^15^. We performed myeloid-specific *Trem2* deletion (*Trem2*^ΔCX3CR1^) specifically during the regression period by tamoxifen treatment, allowing for the assessment of TREM2 function in regression without affecting the progression phase (Figure 1A). This model deletes TREM2 in monocytes and plaque macrophages, as we previously validated^13^. Regressing mice were compared to a baseline group which was euthanized prior to beginning the regression phase.

**Figure 1.**
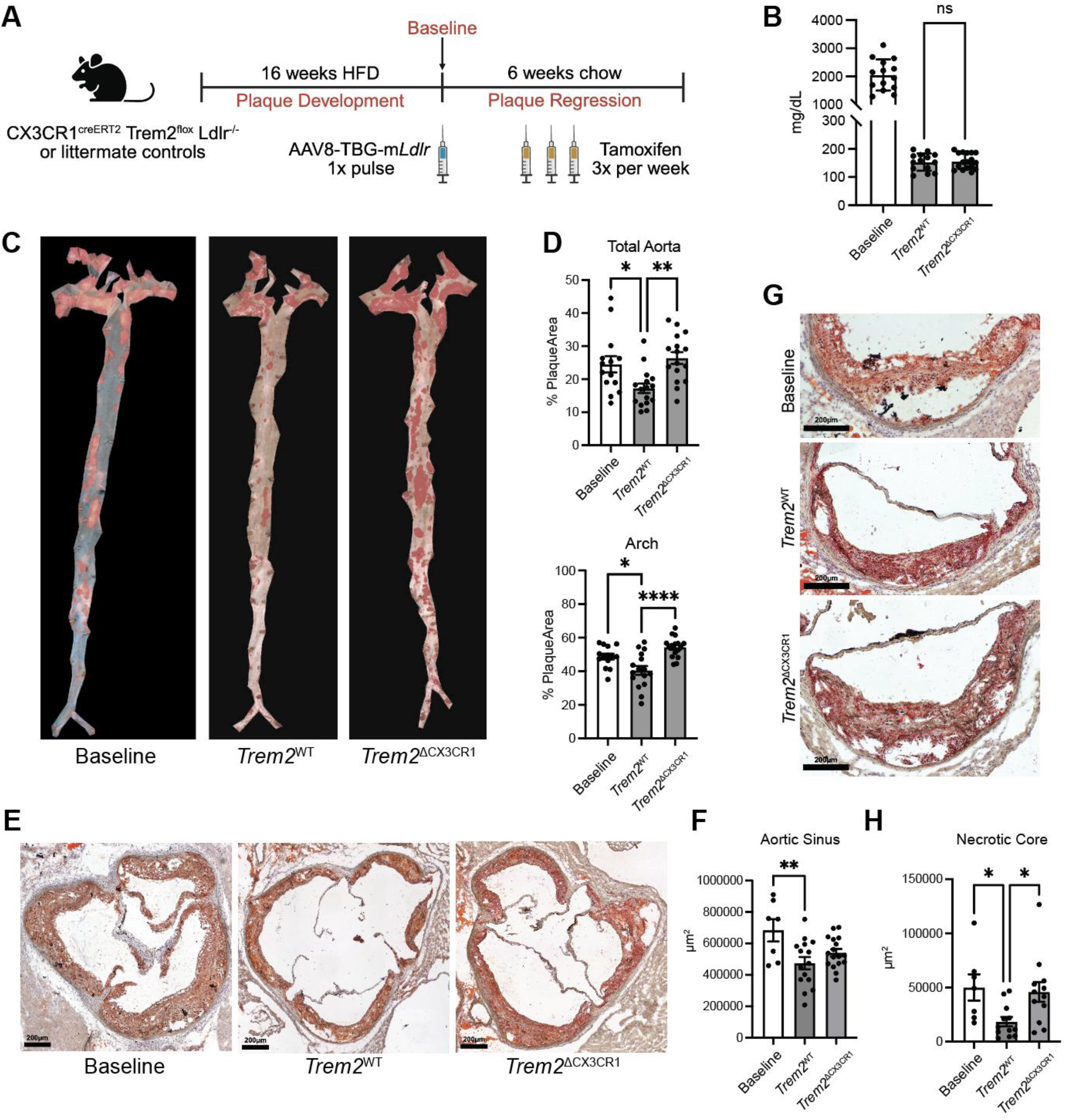
Macrophages require TREM2 for pro-resolving functions during lipid lowering in atherosclerotic plaque. A) Experimental schematic for myeloid-specific *Trem2* deletion regression model. CX3CR1^creER^ *Trem2*^flox^ *Ldlr*^-/-^ mice (*Trem2*^ΔCX3CR1^) or littermate controls were placed on 16 weeks of high fat diet (HFD) to develop plaque. A single injection of a liver-targeted AAV was administered to express *Ldlr*. Mice were then switched to chow diet for the remainder of the study. During the regression period, mice received tamoxifen via oral gavage three times weekly to induce *Trem2* deletion in CX3CR1-expressing cells. B) Quantification of serum cholesterol levels 9 days post-AAV administration. C) *En face* Oil Red O (ORO)-stained images of atherosclerotic aortas from Baseline, *Trem2*^WT,^ and *Trem2*^ΔCX3CR1^ mice. D) Quantification of plaque area. E) ORO-stained images of aortic sinus plaques and F) quantification of plaque area. G) ORO-stained images showing necrotic cores, which are defined as acellular areas within plaque and H) quantification of necrotic core area.

Compared to baseline, there was a 10-fold decrease in serum cholesterol levels in regressing mice, with no genotype effect (Figure 1B). TREM2-sufficient mice had a significant reduction in plaque area compared to the baseline cohort, whereas this was abrogated in *Trem2*^ΔCX3CR1^ animals (Figure 1C-F, Figure S1A-C). Deletion of *Trem2* also impeded the resolution of necrotic cores in aortic sinus plaques during regression (Figure 1G-H), while having no significant impact on circulating immune cells or body weight (Figure S1D-F). Compared to the baseline group, regressing plaques from control mice had reduced macrophage and lipid area but larger smooth muscle actin area within the fibrous cap (Figure 2A-C), a hallmark of plaque stabilization^1,16^. Compared to TREM2-sufficient regressing plaques, *Trem2*^ΔCX3CR1^ plaques had fewer macrophages (Figure 2A-B), which is consistent with a potential role of TREM2 in mediating the efferocytosis-induced proliferative burst^17^ as well as plaque macrophage survival^13^.

**Figure 2.**
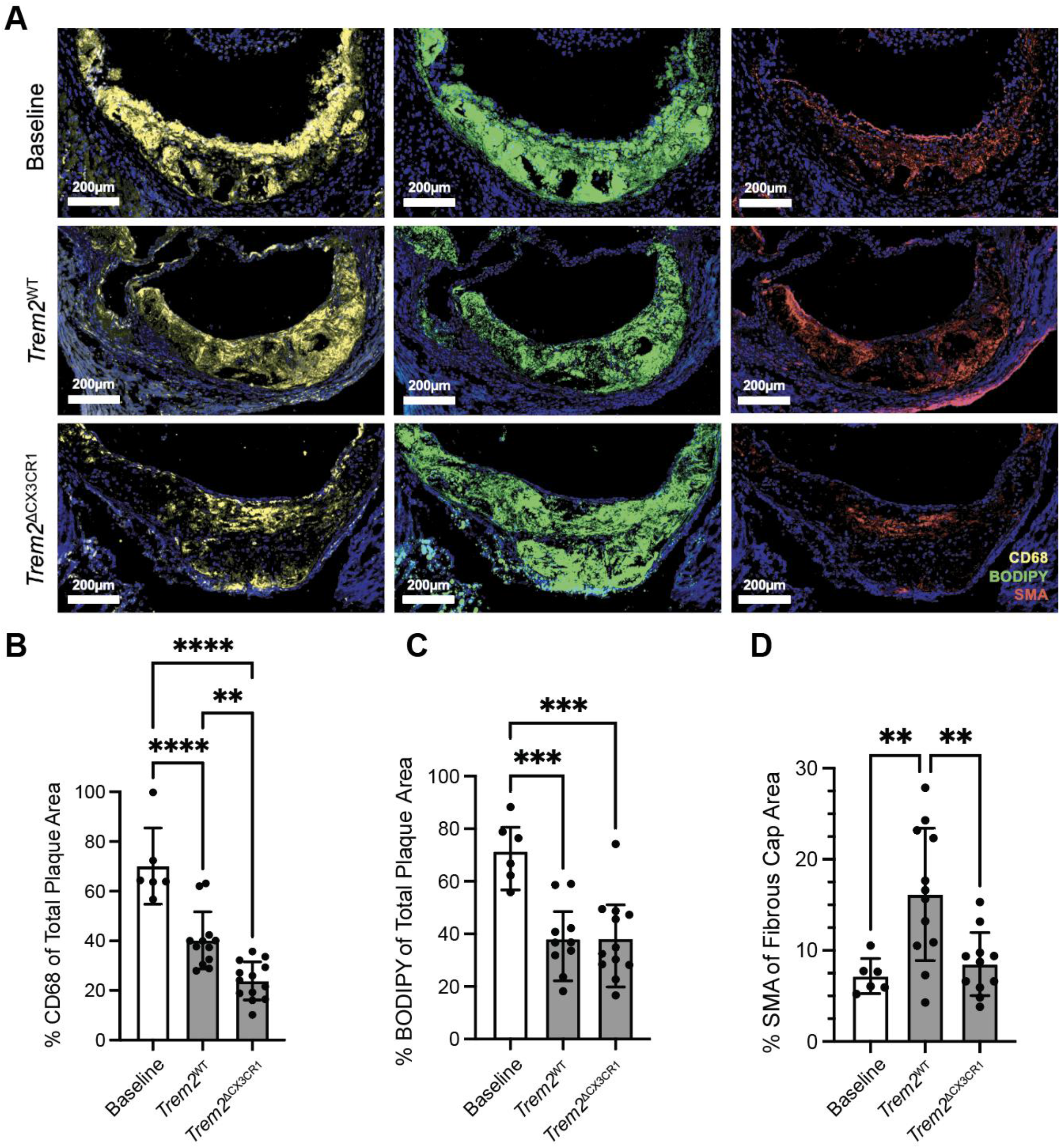
CX3CR1-specific *Trem2* deletion during regression hinders plaque macrophage abundance and fibrous cap formation. A) Immunofluorescence staining macrophages (CD68), lipid content (BODIPY), and smooth muscle actin (SMA) in aortic sinus plaque sections. Images were acquired using confocal microscopy. Associated quantification of B) CD68+ area, C) BODIPY+ area, and D) SMA+ area, expressed as percentage of total plaque area. Fibrous caps were considered as the first ∼50µm from the luminal space.

The contribution of recruited monocytes and plaque resident macrophages in atherosclerosis regression remains controversial^2,3^. Since CX3CR1 is expressed by both cell subsets, we also utilized a CCR2-specific model (*Ccr2^creER^ Trem2^flox^ Ldlr^-/-^, Trem2*^ΔCCR2^) to isolate the impact of TREM2 expression on recruited monocytes during regression. CCR2 is expressed by classical monocytes, which will mature into nonclassical monocytes in the blood^18^. Thus, this model efficiently deletes in all blood monocytes and their progeny, without affecting the existing plaque-associated macrophages that are CCR2-negative. In this monocyte-specific regression model *Trem2*^ΔCCR2^, mice were fed a HFD for 16 weeks, then AAV-*mLdlr* was used to induce regression and *Trem2* deletion was induced by providing TAM by gavage (Figure 3A). *Trem2*-sufficient and -deficient mice regressed to a similar degree, where *Trem2* deficiency on recruited monocytes did not impact plaque area, macrophage and lipid area, or stability features (Figure 3-4). There was also no difference in circulating immune cells or body weight between *Trem2^WT^* and *Trem2*^Δ*CCR2*^ mice during regression (Figure S2). Fate mapping studies using genetic labeling of recruited monocytes throughout the 6 week regression period (*Ccr2^creER^ R26^tdTomato^ Ldlr^-/-^)* showed that monocytes recruited into lesions failed to penetrate beyond the plaque surface and were unable to localize near the necrotic cores, which is consistent with prior studies using a monocyte bead labeling approach^2,15^ (Figure 5A-B). Importantly, *Trem2* deletion did not impact the monocytes’ ability to enter the plaque, as the number of labeled cells recruited during the regression phase remained comparable between genotypes (Figure 5C). These data support that atherosclerotic plaque resolution relies on *Trem2*-dependent phenotype switching of plaque macrophages, not monocyte-derived cells, to mediate regression following cholesterol lowering.

**Figure 3.**
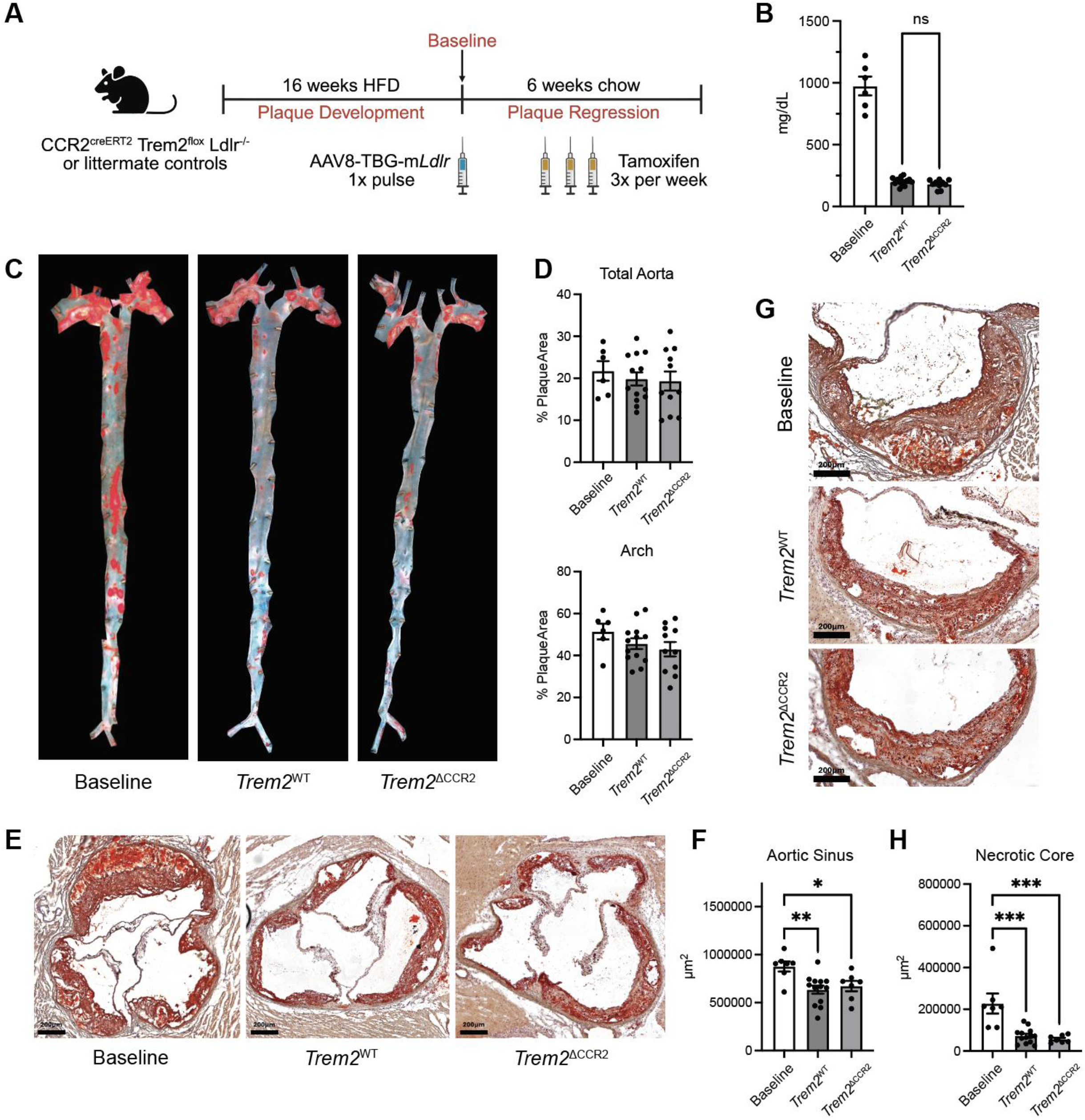
*Trem2* deletion in newly recruited monocytes during atherosclerosis regression does not impact plaque resolution. A) Experimental schematic for monocyte-specific *Trem2* deletion regression model. CCR2^creER^ *Trem2*^flox^ *Ldlr*^-/-^ mice (*Trem2*^ΔCCR2^) or littermate controls were placed on 16 weeks of high fat diet (HFD) to develop plaque. A single injection of a liver-targeted AAV was administered to express *Ldlr*. Mice were then switched to chow diet for the remainder of the study. During the regression period, mice received tamoxifen via oral gavage three times weekly to induce *Trem2* deletion in CCR2-expressing cells. B) Quantification of serum cholesterol levels 9 days post-AAV administration. C) *En face* Oil Red O (ORO)-stained images of atherosclerotic aortas from Baseline, *Trem2*^WT^, and *Trem2*^ΔCCR2^ mice. D) Quantification of plaque area. E) ORO-stained images of aortic sinus plaques and F) quantification of plaque area. G) ORO-stained images of necrotic core area in aortic sinus plaques and H) quantification of necrotic core area.

**Figure 4.**
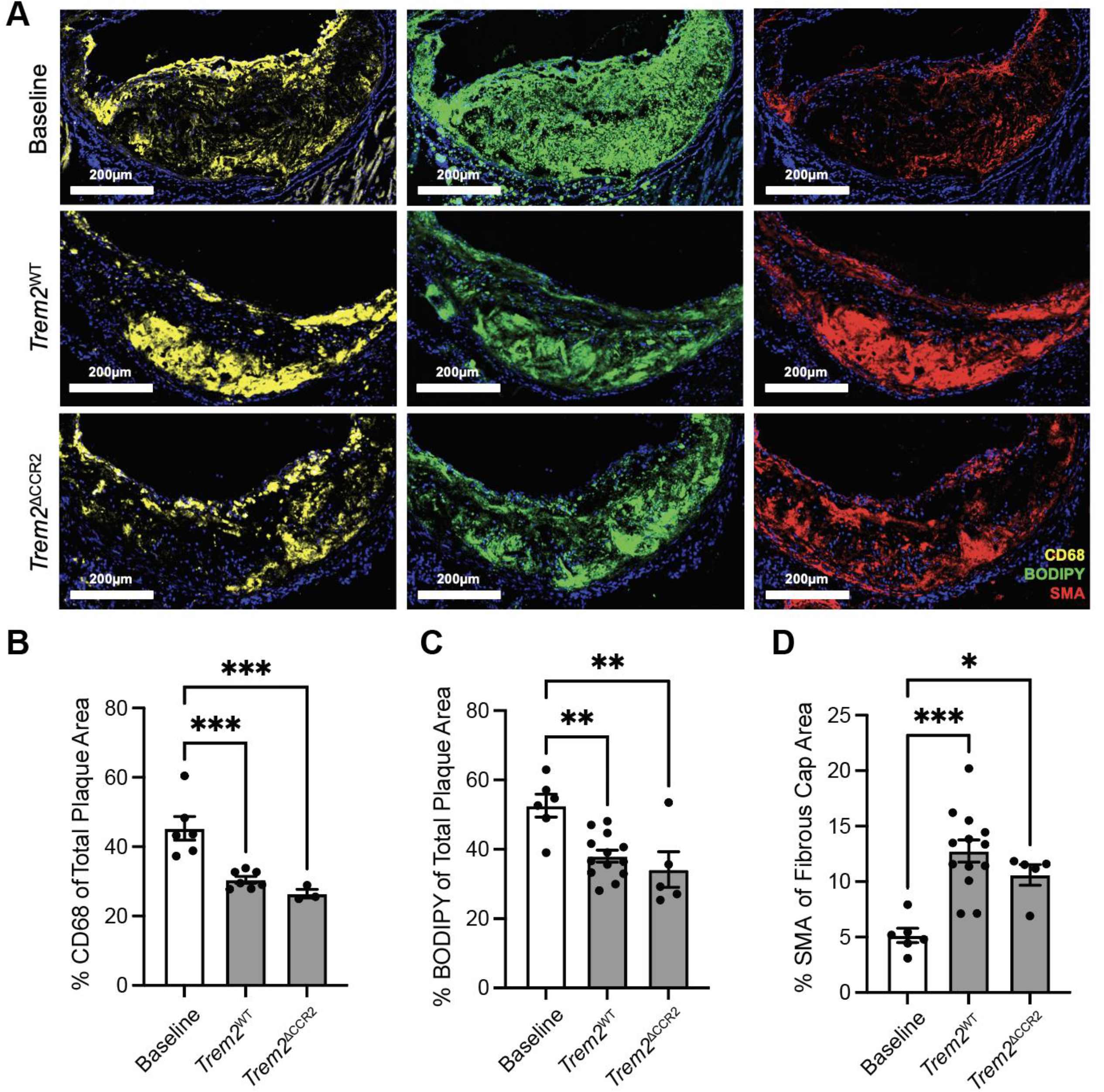
CCR2-specific *Trem2* deletion does not impact plaque macrophage abundance or fibrous cap formation during regression. A) Immunofluorescence staining macrophages (CD68), lipid content (BODIPY), and smooth muscle actin (SMA) in aortic sinus plaque sections. Images were acquired using confocal microscopy. Associated quantification of B) CD68+ area, C) BODIPY+ area, and D) SMA+ area, expressed as percentage of total plaque area. Fibrous caps were considered as the first ∼50µm from the luminal space.

**Figure 5.**
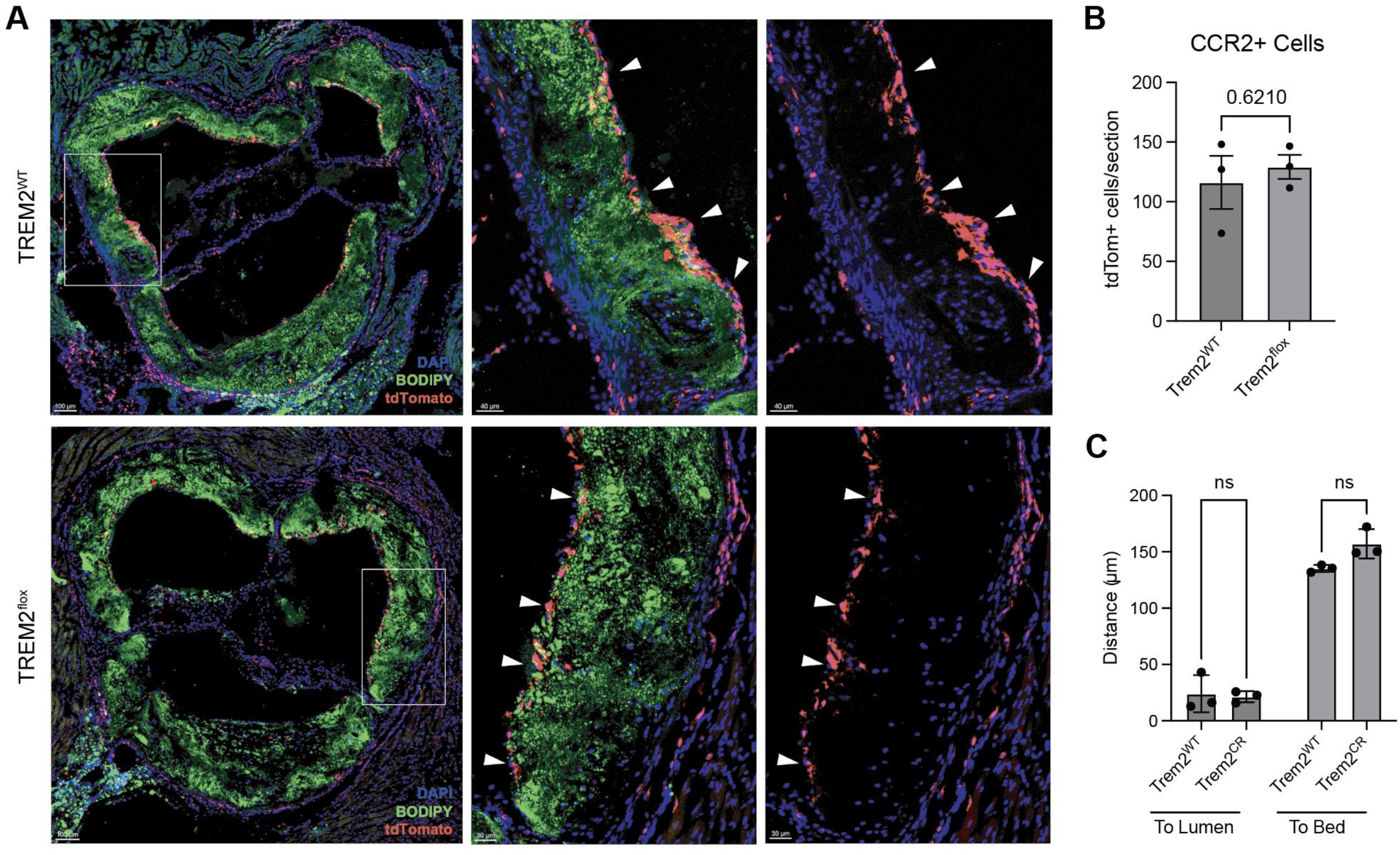
Monocytes recruited to plaque during regression are spatially restricted from necrotic cores. A) CCR2^creER^ *Trem2*^flox^ R26^tdTomato^ *Ldlr*^-/-^ mice or control were administered tamoxifen three times per week by oral gavage during the regression period only to label any CCR2-expressing cells recruited to the plaque. DAPI (blue), BODIPY (green), and tdTomato (red) staining of aortic sinus sections was visualized by confocal microscopy. B) Quantification of the distance of each tdTomato+ cell to either the artery lumen or the artery bed. Three plaque sections for each mouse were averaged. C) Quantification of tdTomato+ cells per section. Three plaque sections for each mouse were averaged.

### Ex-Lipid associated macrophages (ExLAMs) have enhanced efferocytic capacity

Given these *in vivo* results, we sought to understand how TREM2-expressing lipid-associated plaque macrophages change in response to lipid unloading. Efferocytosis, or the clearance of dead cells and debris, is a core homeostatic macrophage function in which TREM2 plays a known role^5,6,13,14^. This also represents a possible mechanism for the TREM2-dependent necrotic core resolution observed in the *in vivo* regression model. The impact of cholesterol removal from LAMs, and its influence on macrophage plasticity and function, remains unexplored.

To determine how LAM function is altered following cholesterol efflux, bone marrow-derived macrophages (BMDMs) were loaded with cholesterol overnight to generate LAMs, followed by two days of cholesterol efflux to generate ExLAMs (Figure 6A). Unloading of lipids was confirmed by BODIPY neutral lipid staining and mean side scatter by flow cytometry (Figure 6B-C). ExLAMs also maintained high expression of LAM- and efferocytosis-associated proteins TREM2, CD9, CD36, and MerTK, even though they were no longer exposed to cholesterol and had cleared intracellular lipid droplets (Figure S3A). ExLAMs did not die in response to lipid unloading (Figure S3B) but underwent a proliferative burst (Figure S3C-D), as measured by BrdU incorporation.

**Figure 6.**
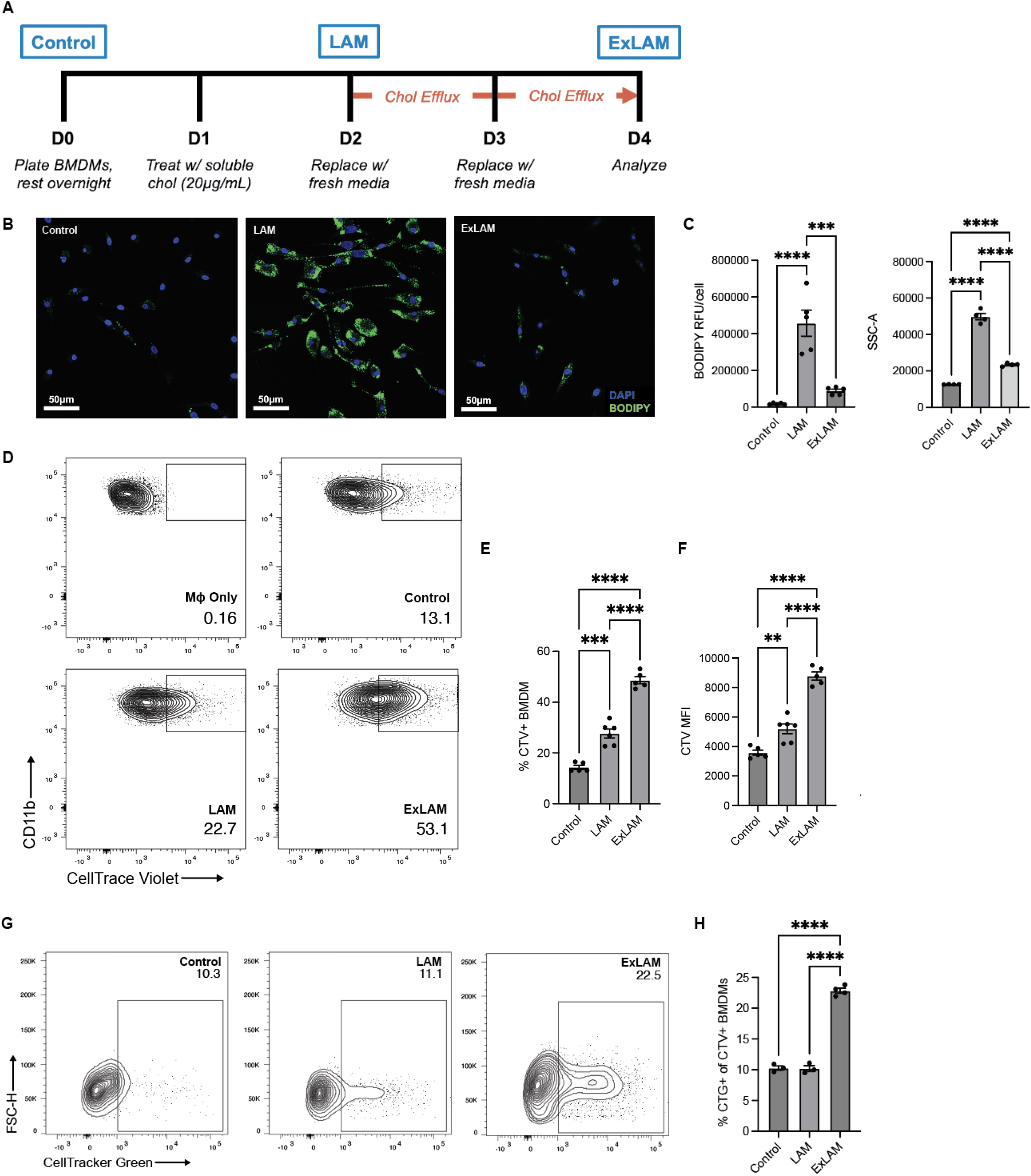
Lipid unloaded ex lipid associated macrophages (ExLAMs) have enhanced efferocytic capacity. A) Experimental setup for ExLAM cell generation. Briefly, BMDMs were loaded with 20 µg/mL soluble cholesterol overnight to become LAMs and then allowed to efflux cholesterol over a two-day period before being used in assays. LAMs (20 µg/mL soluble cholesterol overnight) and Control (no cholesterol). B) Immunofluorescent imaging of bone marrow-derived macrophages (BMDMs) stained for nuclei (DAPI) and lipid content (BODIPY). C) Quantification of BMDM lipid content (BODIPY) via imaging and side scatter area (SSC-A) via flow cytometry. D) Representative contour flow cytometry plots showing BMDMs (CD11b+ F4/80+) that have engulfed a CellTrace Violet (CTV)-labeled apoptotic splenocyte. E) Quantification of (D) by flow cytometry. F) Mean fluorescent intensity (MFI) of TREM2 expression on efferocytes from (D). G) Representative contour flow cytometry plots showing BMDMs that have engulfed a CellTracker Green (CTG)-labeled apoptotic splenocyte after engulfing a CellTrace Violet (CTV)-labeled splenocyte. Cell are pregated on CTV+ BMDMs, as shown in Figure S3A. H) Quantification of (G).

Assessment of efferocytic capacity of untreated control, LAM, and ExLAM macrophages with irradiated CellTrace Violet (CTV)-labeled apoptotic cells (ACs)^13^ revealed that ExLAMs had significantly higher engulfment of ACs compared to control macrophages or LAMs (Figure 6D-E, S3E-F), providing evidence of LAM plasticity and enhanced effector function following cholesterol efflux. ExLAMs also showed elevated TREM2 expression when exposed to ACs (Figure 6F, S3G). In a continuous efferocytosis assay, ExLAMs showed enhanced efferocytic capacity to engulf multiple apoptotic cells sequentially. BMDMs were co-cultured with multiple rounds of distinctly labeled ACs and analyzed for engulfment of multiple ACs. While LAMs uptake more single ACs than control macrophages, both populations showed equivalent uptake of a second AC. However, ExLAMs exhibited enhanced primary uptake and secondary AC uptake at more than twice the frequency of LAMs and controls (Figure 6G-H). Together, these results suggest that clearance of lipids from LAMs, generate functionally distinct cells, ExLAMs, that possess enhanced proliferation and efferocytosic function. This may have implications in limiting secondary necrosis in advanced atherosclerotic plaques, where dead cells far outnumber functional live macrophages.

### Lipid unloading of ExLAMs drives distinct transcriptional changes while maintaining core LAM signature

Next, we explored the transcriptional profile of ExLAMs by bulk RNA sequencing. ExLAMs and LAMs express similarly high numbers of differentially expressed genes (DEGs) compared to controls, with a significant number of those genes being shared between LAMs and ExLAMs (Figure 7A). Interestingly, ExLAMs cluster independently of both control BMDMs or LAMs by principal component analysis (Figure 7B), supporting that once lipid droplets have been cleared, ExLAMs do not transcriptionally revert to the untreated condition. Applying a canonical LAM gene signature^19^ showed that ExLAMs maintain upregulation of *Cd63, Fabp5,* and *Lgals3* during lipid unloading and further upregulated *Pld3, Cd68, Ctsb,* and *Ctsd* compared to the LAM state (Figure 7C). Many of these genes are associated with lysosomal degradation, a key component of debris clearance. Similarly, ExLAMs upregulate numerous efferocytosis genes, with *Itgb3, Dock1, Stab2, Mfge8,* and *Mertk* among the highest expressed (Figure 7D). Compared to controls, the top DEGs in ExLAMs include those involved in cholesterol efflux (*Abca1, Abcg1*), cytoskeletal organization (*Itgb3, Itgb2, Timp2, Adamtsl5*), and survival (*Cd5l, Gpx3*). Top DEGs in the control condition include those involved in type 1 interferon response (*Oas1g, Oasl2, Irf7*) and activation (*Ccr2, Cd74, Ctnnd2, C3*) (Figure 7E-F). Compared to ExLAMs, LAMs upregulated genes associated with NFκB activation (*Nfkbia, Nfkb2, Nfkbie, Relb*) (Figure S5A-B). Altogether, transcriptional changes in ExLAMs suggest that they retain a key gene program associated with LAM formation, but upregulate lysosomal, extracellular matrix, and cytoskeletal remodeling genes to support the enhanced efferocytic and proliferative capacity observed in functional assays.

**Figure 7.**
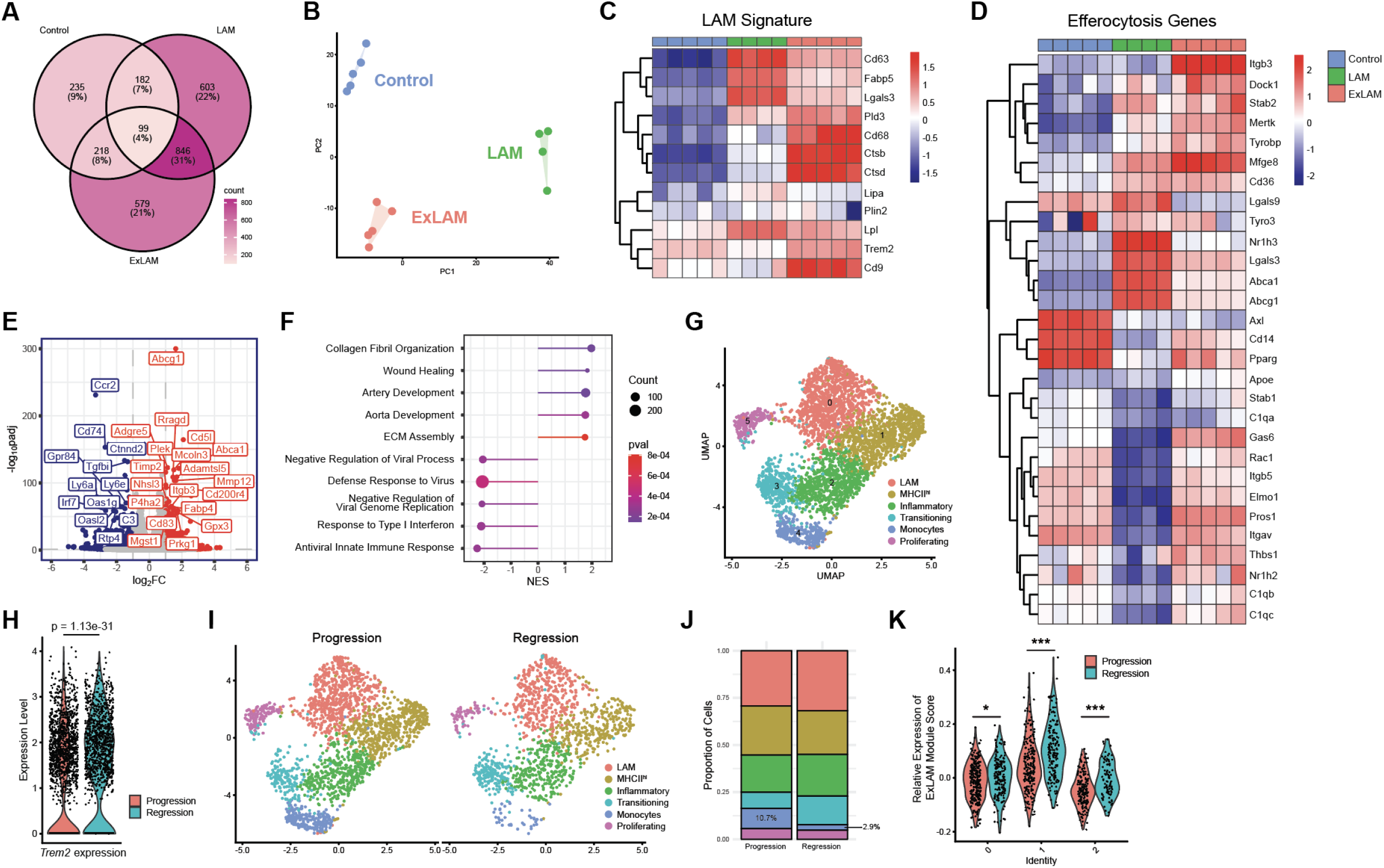
Lipid unloading induces transcriptional changes associated with engulfment and lysosomal degradation while maintaining LAM signature. A) Venn diagram showing the number of differentially-expressed genes (DEGs) in control, LAMs, and ExLAMs. B) Principal component analysis (PCA) plot of bulk RNA sequencing data of mouse BMDMs. Top 2000 genes were considered. C) Heat map of relative expression of lipid-associated macrophage genes across the three treatment groups. Data shown are row-wise Z scores. D) Heat map of relative expression of efferocytosis-associated genes across the three treatment groups. Data shown are row-wise Z scores. E) Volcano plot of differentially expressed genes between ExLAMs and control BMDMs. padj <0.05 and abs(Log_2_FC) > 1 was considered significant. Genes in red are significantly upregulated in ExLAMs and genes in blue are significantly downregulated. F) Pathway analysis using DEGs from (E). Gene Ontology Biological Processes (GOBP) pathway library was utilized. G) Uniform Manifold Approximation and Projection (UMAP) clustering of sorted CD11b+ CX3CR1+ cells from a total atherosclerotic aorta digest. Dataset was generated by Lin et al and reanalyzed independently. Adventitial macrophages were excluded based on high expression of *Lyve1*. Cell clusters were named using SingleR and manual identification. H) *Trem2* expression on all clusters between experimental groups. I) Clusters from (G) split by experimental group and J) cluster proportions split by experimental group. K) An “ExLAM Module Score” was generated using the top 50 upregulated genes from ExLAM bulk RNA sequencing. This score was then projected onto the subclusters from Figure S5G and compared between the two experimental groups.

We also investigated the impact of *Trem2* deletion (*Trem2^-/-^)* on ExLAM transcriptional programming *in vitro*. In comparison to WT ExLAMs, *Trem2*^-/-^ ExLAMs failed to upregulate a subset of LAM- and efferocytosis-associated genes when assessed by bulk RNA sequencing (Figure S6A-F). In support, expression of the Apoptotic Cell Clearance pathway (GOBP) was diminished while Negative Regulation of Wound Healing (GOBP) was enriched (Figure S6G). Similar to the role of *Trem2* in programming foam cell formation, these data further suggest that *Trem2* plays a key role in these macrophage pro-resolving processes and support the prior *in vivo* deletion results^13^.

An ExLAM-like population with shared transcriptional features could also be found in an *in vivo* setting of lipid lowering. Comparison of a single cell RNA sequencing dataset of murine aortic plaque macrophages during atherosclerosis progression with hypercholesterolemia and atherosclerosis regression, where hypercholesterolemia was reversed for 2 weeks^20^ identified six clusters of monocyte and macrophages based on the expression of canonical genes (Figure 7G, Figure S7A-C). Overall, higher expression of *Trem2* was found across all clusters (Figure 7H, Figure S7D). When split between treatment conditions, the regression group showed fewer total macrophages in the samples, which was most clearly associated with a loss of the monocyte cluster compared to the progression sample (Figure 7I-J). *Trem2* expression was associated with the transcriptional transition of monocytes to mature macrophages in both progression and regression settings, as predicted by pseudotime analysis (Figure S7E). The mature foam cell (or LAM) cluster 0 also showed upregulation of several genes involved in membrane dynamics (*Itgb1, Iqgap1, Myof, Plec*), lysosomal degradation (*Lyz1, Lyz2, Ctsb*), efferocytosis (*Lrp1*), and anti-inflammatory polarization (*Mrc1, Ncl*) in regression compared to progression, further supporting the *in vitro* findings in ExLAMs (Figure S7F). Because ExLAMs share gene expression with LAMs, the LAM population was further resolved into three subclusters (Figure S7G-H). By applying an “ExLAM Score” comprised of the top 50 upregulated genes from *in vitro*-generated ExLAMs, this showed the ExLAM associated genes were highest in subcluster 1, the only subcluster that was also more abundant during regression compared to progression. Moreover, the ExLAM score was further upregulated in regression compared to progression (Figure 7K, S7I). Together, these data support that plaque associated foam cells mimic reprogramming events observed in the in vitro ExLAM experiments.

Finally, to test whether ExLAMs are also found in other tissue sites, isolated peritoneal macrophages (PMACs) from HFD-fed and atherosclerosis regression mice were examined by RNA-seq. Indeed, PMACs from regression mice unload lipid similarly to BMDM-derived ExLAMs and showed a strikingly similar DEG profile to the BMDM-derived ExLAMs (Figure S8). Overall, these functional and transcriptional data support that in vivo generated ExLAMs maintain expression of LAM genes, including *Trem2*, while upregulating genes associated with apoptotic cell engulfment and degradation.

### Enhancing TREM2 Signaling Boosts Atherosclerosis Regression and Macrophage Efferocytosis

Our data suggests TREM2 is a key driver of atherosclerosis regression and ExLAM programming. To interrogate the translational potential of TREM2 as a therapeutic target to control pro-resolving macrophage function, we used two approaches to enhance TREM2 signaling in the context of lipid unloading. First, a monoclonal TREM2 antibody agonist (AL002a, Alector Therapeutics) was used in an atherosclerosis regression model. This antibody treatment was previously shown to modulate plaque macrophage function in atherosclerosis progression, mediating foam cell proliferation and enhancing features of plaque stability^11^. TREM2 agonist was provided only during the regression phase, and mice were sacrificed after 4 weeks of regression to allow capture of any accelerated regression features (Figure 8A). Compared to isotype control treated mice, aortae from mice receiving TREM2 agonist showed accelerated plaque regression in the thoracic and abdominal regions, as well as the total plaque area (Figure 8B-C), aligning with the findings in the inducible deletion model. Plaque area and necrotic core area in the aortic sinus of the heart showed similar levels of clearance to the control group (Figure 8D-E).

**Figure 8.**
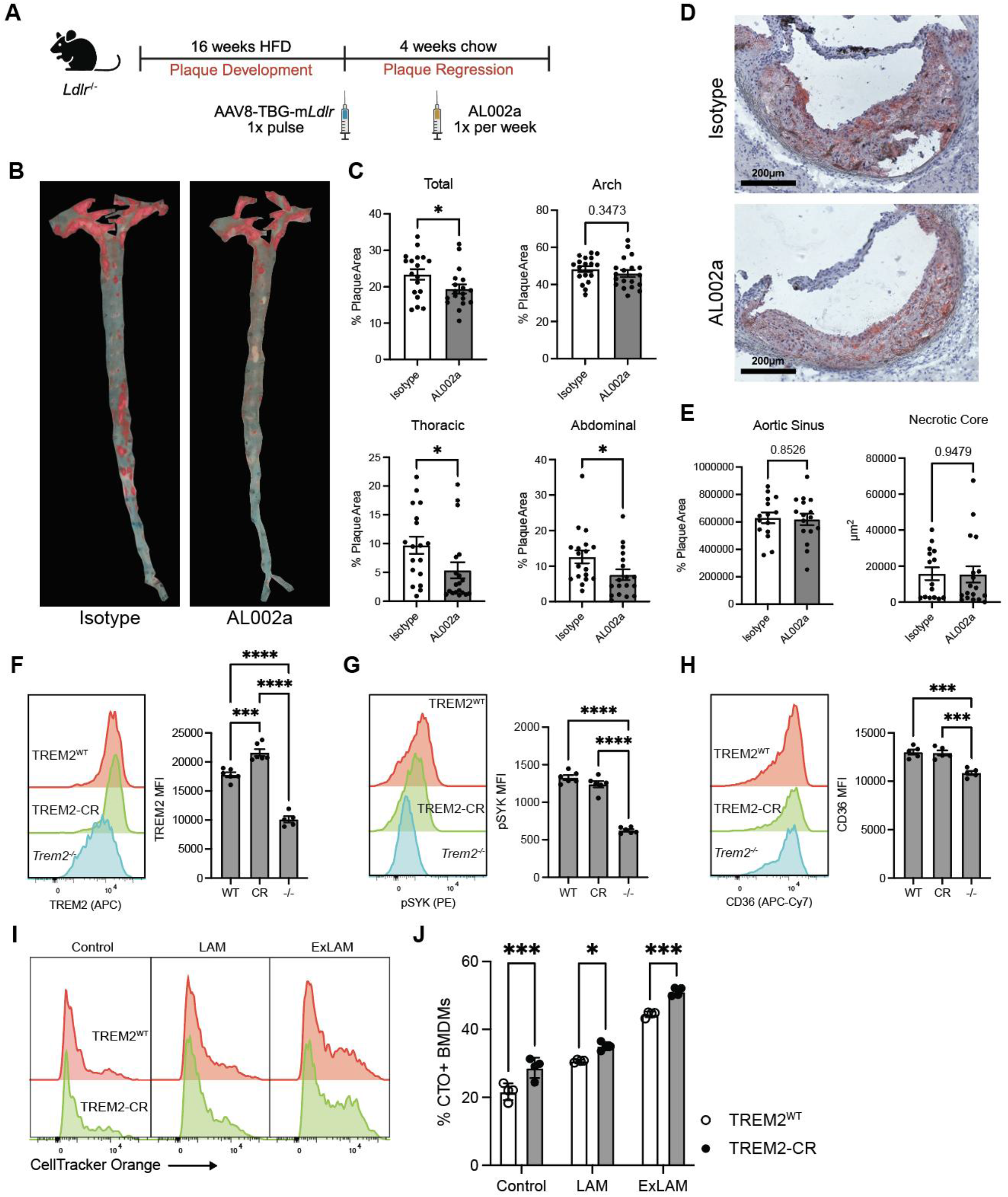
Enhanced TREM2 signaling boosts macrophage efferocytosis and pro-resolving function. A) Experimental schematic for TREM2 agonist regression model. *Ldlr^-^*^/-^ mice were placed on 16 weeks of high fat diet (HFD) to develop plaque. A single pulse of a liver-targeted AAV was administered to re-express *Ldlr*. Mice are then switched to chow diet for the remainder of the study. During the regression period, mice received TREM2 agonist (AL002a) via intraperitoneal injection once per week to stimulate TREM2 signaling. B) *En face* Oil Red O (ORO)-stained images of atherosclerotic aortas from isotype- and AL002a-treated mice. C) Quantification of plaque area. D) Representative ORO-stained images of aortic sinus plaques and E) quantification of aortic sinus plaque area and necrotic core area. F) Flow cytometry histograms and quantification of the mean fluorescent intensity of TREM2, G) phosphorylated SYK (pSYK), and H) CD36 in wildtype (WT), TREM2 cleavage resistant (TREM2-CR), and *Trem2*^-/-^ (KO) bone marrow-derived macrophages loaded with 20 µg/mL of soluble cholesterol overnight. I) Fluorescently labeled WT and TREM2-CR BMDMs were co-cultured and exposed to CellTrace Orange/CTO-labeled apoptotic cells. Uptake of CTO+ apoptotic cells was measured by flow cytometry. J) Quantification of (I).

In a second model of enhancing TREM2 function, a cleavage-resistant TREM2 mouse model (TREM2-CR) was used to stabilize surface expression levels. TREM2-CR mice possess a mutant TREM2 transgene where the native stalk region was replaced with the CD8 stalk region, eliminating the cleavage site. TREM2 proteolytic cleavage is mediated by members of the disintegrin and metalloproteinase domain-containing protein (ADAM) family in the stalk region of the receptor and is often cleaved in inflammatory settings including advanced atherosclerosis^14,21^. The shed extracellular domain of the receptor (sTREM2) can be detected in blood and cerebrospinal fluid and is correlated with disease severity in both Alzheimer’s and cardiovascular disease^14,22^.

TREM2-CR macrophages effectively stabilized TREM2 protein on the cell surface, showing higher TREM2 protein levels after lipid loading and maintaining normal intracellular signaling of pSYK (Figure 8F-G). CD36, a molecule previously linked to TREM2 expression^19,23^ was found to be expressed at a normal level in lipid loaded TREM2-CR BMDMs (Figure 8H). To test whether enhanced TREM2 expression could mediate beneficial effects in ExLAMs, TREM2^WT^ and TREM2-CR BMDMs were co-cultured to test head-to-head efferocytic capabilities of these cell lines under control, LAM, or ExLAM conditions. TREM2-CR BMDMs showed enhanced efferocytosis compared to TREM2^WT^, and this was true in each of the control, LAM, and ExLAM conditions (Figure 8I-J). Together, these data suggest that modifying TREM2 through enhanced signaling by antibody agonist, or through receptor stabilization on the cell surface, are effective approaches for stimulating macrophage pro-resolving functions in atherosclerosis regression and driving the function of ExLAMs.

## DISCUSSION

This study identifies and characterizes a previously unrecognized mechanism for TREM2-dependent LAM reprogramming and the sustained functional effects on atherosclerotic plaque remodeling. Using a combination of *in vivo* loss-of-function models, functional assays, and transcriptomic analysis, we show that TREM2 signaling on plaque macrophages, but not newly recruited monocytes, drives plaque regression and necrotic core resolution. We propose that lipid lowering generates the formation of lipid experienced ExLAMs. ExLAMs have a distinct transcriptomic profile associated with prior lipid exposure that is preserved following lipid unloading and promotes enhanced engulfment, lysosomal degradation, and matrix remodeling, which was inhibited by *Trem2* deletion. This suggests that lipid-driven macrophage phenotype switching may be harnessed through TREM2 to drive inflammation resolution in atherosclerotic disease. Notably, therapeutic targeting of TREM2 through antibody agonism and cleavage resistance also boosted atherosclerosis regression and efferocytosis. This finding unveils TREM2 as a viable therapeutic target for modulating LAM reprogramming and plaque resolution following aggressive lipid lowering therapy in cardiovascular disease patients.

These findings challenge the prevailing theory that the plaque associated foam cells or LAM are a terminal, dysfunctional macrophage state. Moreover, it challenges the paradigm that atherosclerosis regression is mediated by newly recruited monocytes^3^. Instead, our data support that lipid lowering leads to long-term enhancement of macrophage efferocytic function in preexisting lipid-loaded cells and drives beneficial plaque remodeling. While prior studies have associated the presence of LAMs with disease severity in many contexts^8^, the possible fates of LAMs and the ability to reprogram their function has not been systematically interrogated until now. Moreover, we position TREM2 as a leading target for leveraging LAM-to-ExLAM rejuvenation. Our work ultimately frames lipid loading as a dynamic programming event that shapes macrophage function, not merely a consequence of inflammation or disease.

This revised model has implications that extend beyond cardiovascular disease to other contexts in which macrophages encounter lipid-rich environments, including obesity, neurodegeneration, chronic infection, and cancer. In these settings, the capacity of macrophages to resolve lipid loading may represent a general mechanism linking macrophage metabolic adaptation to the restoration of tissue homeostasis and repair across diverse conditions. In obesity and metabolic disease, GLP-1 agonists modulate gastrointestinal metabolism to drive insulin sensitivity, reduce glucose output, and enhance β cell survival, leading to dramatic weight loss in many individuals^24,25^. In these systems, we hypothesize that ExLAMs will form and potentially play integral roles in tissue remodeling and disease reversal. In addition, targeting TREM2 in combination with these treatments may lead to accelerated and improved outcomes for the established therapeutics.

While weight loss generally promotes cardiovascular health, studies show that repeated weight cycling disrupts adipose tissue homeostasis and sustains inflammation and cardiovascular disease risk^26–28^. Similar observations have been made in cardiovascular disease models following repeated HFD-feeding exposure^29,30^. Future work will need to refine the cellular dynamics of repeated lipid challenge and the response of LAM/ExLAM cells in atherosclerotic disease and other organs. Finally, in numerous cancer types, LAMs are present and are typically associated with an immunosuppressive tumor microenvironment^31–33^. Existing studies demonstrate that blocking lipid uptake and storage via scavenger receptor blockade potently reduced lipid load and tumor growth^34,35^, but further studies are needed to determine whether rewiring LAM metabolism and function through lipid unloading can provide similar immune boosting effects in a cancer model. In all, understanding how macrophages respond to altered lipid burden, and the long-term programming that is mediated by lipid loading and unloading, will uncover fundamental mechanisms that govern their functional plasticity and reveal new therapeutic opportunities to restore tissue homeostasis in numerous human diseases.

## METHODS

### Mice

All experiments and procedures were approved by the University of Minnesota (UMN) Institutional Animal Care and Use Committee (IACUC) and the UMN Institutional Biosafety Committee (IBC). Mouse strains used for this study include: B6 (C57BL/6, Jackson Laboratory (JAX), 000664); *Ldlr^−/−^* (B6.129 S7-Ldlr^tm1Her^/J, JAX, 002207); Cx3cr1^creER^ (B6.129P2(C)-Cx3cr1^tm2.1(cre/ERT2)Jung^/J, JAX, 020940); *Ccr2^creER^ (*^36^); *Trem2^−/−^* (developed and provided by Marco Colonna, Washington University in St. Louis), *Trem2^flox^* (B6(C3)-TREM2^tm1c(EUCOMM)Wtsi^/AdiujJ, developed and provided by Bruce Lamb at Indiana University, JAX, 029853); *R26^tdTomato^*(B6.Cg-Gt(ROSA)26Sor^tm9(CAG-tdTomato)Hze^/J, JAX, 007909), and TREM2-CR (TREM2 Cleavage Resistant) (developed and provided by Marco Colonna, Washington University in St. Louis). All mice are on the C57BL/6 background and bred in specific pathogen-free animal facilities maintained by UMN Research Animal Resources. When possible, littermates were used for experiments. All mice were enrolled in the study between 6 to 12 weeks of age. Facilities were maintained at ∼23 °C with a 12-h light/dark cycle. Cages were changed weekly, and water was freely available through a Lixit valve. All experiments were performed in both male and female mice.

### HFD Feeding

All animals in this study were fed *ad libitum.* High fat diet (HFD) (Envigo Teklad #TD.88137), an adjusted calorie diet containing 42% fat, was fed to mice during plaque development. Littermate control animals (baseline groups) were fed HFD for 16 weeks only. Regression mice were fed HFD for 16 weeks before switching to chow for 6 weeks subsequently following AAV administration.

### AAV Administration

To introduce expression of *Ldlr* in hepatocytes of *Ldlr*^-/-^ mice, we employed an adeno-associated virus (AAV) vector to deliver the mouse *Ldlr* gene. AAV8-TBG-m*Ldlr* was purchased from Vector Biolabs (Lot #241104-20-23). Liver specificity of the vector is conferred by the AAV serotype 8’s tropism for the liver in combination with the thyroxine binding globulin (TBG) promoter, which restricts expression to hepatocytes. Baseline control mice were euthanized prior to AAV administration. For mice continuing in the regression study, a single pulse of AAV was administered at a dose of 2.5x10^12^ GC/mouse via tail vein injection. Blood was collected at 9 days post-AAV and again at the end of the study to confirm lowering of serum cholesterol levels using the Cholesterol E measurement kit (FujiFilm #999-02601). Subjects with serum cholesterol levels above 200 mg/dL at either time point were excluded from the study. Regression mice were switched to chow diet immediately following AAV administration until the end of the study.

### Tamoxifen Administration and Plaque Regression

Beginning one day following AAV administration, regression mice began receiving tamoxifen (TAM) via oral gavage to drive CreER-induced deletion of *Trem2*. TAM (Sigma Aldrich #T5648-5G) was dissolved in corn oil at 20 µg/mL. 200 µL per mouse was given three times weekly for the duration of the 6 week regression period. All mice in the regression study were given TAM regardless of genotype.

### Aortic Plaque Analysis

Aortas from mice were harvested and fixed in 4% PFA overnight at 4°C. Perivascular adipose tissue was removed manually under a stereomicroscope, and the vessel was cut along the greater and lesser curvatures, then pinned *en face* to wax dishes. Pinned samples were washed three times with water and incubated in freshly prepared Oil Red O (Sigma Aldrich #00625-25G) solution for 3 hours protected from light at RT. Samples were then incubated in 60% isopropyl alcohol for 5 minutes at RT and washed three times with ddH_2_O. Aortae were imaged using a Leica S9i stereomicroscope. Aorta images were merged using Adobe Photoshop and plaque was quantified using ImageJ software.

### Aortic Sinus Imaging and Immunofluorescence

Hearts from mice were harvested and immediately fresh frozen in OCT compound. 10µm sections were cut on a cryostat and mounted onto slides which were then stored at -80°C until staining. For staining, slides were warmed at RT for 10 minutes and excess OCT was washed off in a 1x PBS bath. Slides were then fixed in 4% PFA for 30 minutes at RT followed by blocking and permeabilizing in a 5% donkey serum in 1% Triton X-100 solution. To quantify plaque area, slides were stained in Oil Red O for 30 minutes and washed 3 times in a 1x PBS bath. Cover slips were mounted using Fluoromount-G (Southern Biotech #0100-01) and allowed to cure overnight in the fridge. Slides were imaged on a Leica SP8 inverted confocal microscope with an attached brightfield light source and quantified using ImageJ. For immunofluorescence, primary antibodies were added at 1:500 and stained at RT for 1 hour, followed by secondary antibodies added at 1:1000 and stained at RT for another 1 hour. The following antibodies were used: CD68 primary (clone FA-11, Biolegend #137002), Actin primary (clone 1A4, EMD Millipore #113200). Slides were washed 3 times in a 1x PBS bath and mounted with Fluoromount-G and allowed to cure overnight in the fridge. Slides were imaged using a Leica SP8 confocal microscope and quantified using QuPath and ImageJ. For both Oil Red O and immunofluorescence, 2-3 sections per mouse were quantified and averaged for data presentation.

### TREM2 Agonist Antibody Treatment

As described above, *Ldlr*^-/-^ mice were fed HFD for 16 weeks continuously before receiving a single pulse of AAV to regain cholesterol control. Mice were then switched to chow diet for the remainder of the study. During the regression period, mice received intraperitoneal injections one time per week of isotype control or TREM2 agonist (AL002a, Alector Therapeutics) at 40 mg/kg for 4 weeks.

### Single Cell RNA Sequencing Analysis

Publicly available scRNA sequencing datasets were downloaded from Gene Expression Omnibus (GEO, GSM3507490 and GSM3507491^20^) and all analysis was performed in R software (version 3.19). Quality control entailed filtering out cells with nFeature_RNA < 500, nCount_RNA > 15000, and mitochondrial gene percentage > 5%. Filtered progression and regression datasets were then merged. The Harmony package was utilized to correct for any potential batch effects. Normalization and scaling were performed using the NormalizeData and ScaleData functions, respectively, within the Seurat package. Principle Component Analysis (PCA) was performed with the RunPCA function, and two-dimensional data visualization was performed using RunUMAP. Both functions utilized the first 20 principal components. Cells clusters were identified using canonical gene markers: foam cells (*Cd9* and *Spp1*), MHCII^hi^ macrophages (expression of many MHC-associated genes including *H2-Aa* and *H2-DMa*), inflammatory macrophages (expression of many cytokines and interferon response genes including *Ccl7* and *Oasl1*), monocytes (*Ms4a4c* and *Zpb1*), and proliferating macrophages (*Top2a* and *Mki67*). Transitioning cells were identified as expressing genes found in both foam cell and inflammatory clusters, albeit at lower levels, as well as their position in the pseudotime between monocytes and more differentiated populations. Full list of cluster-defining genes can be found in the supplemental materials. The adventitia macrophage cluster was filtered out based on high expression of *Lyve1* to isolate plaque macrophages only. Differential gene expression (DEG) analysis was performed using the FindMarkers or FindAllMarkers packages. Pathway analysis was performed using the fgsea package. The Gene Ontology Biological Processes (GOBP) pathways were utilized from the msigdbr package. All volcano plots and dot plots were generated using the ggplot2 package. Heatmaps were generated using the pheatmap package.

### Trajectory Analysis

Monocle3 package (version 1.3.7) was used to predict pseudotime trajectory across myeloid cell clusters. Scaled and normalized gene expression counts and UMAP dimension reduction were input for pseudotime calculation. To plot the trajectory onto the UMAP, a minimum branch length of 4 was used, and use_partition=F was employed to force the trajectory to continue over gaps in the UMAP. The root node was manually chosen as the monocyte cluster. To produce the pseudotime line plot, cells were grouped based on normalized expression of *Trem2* that was either above (*Trem2*^hi^) or below (*Trem2*^lo^) a value of 0.5. A bin width of 0.2 was utilized, and the root node was manually chosen as the monocyte cluster.

### In Vitro Generation of BMDMs

Bone marrow was isolated from the femur and tibia of mice and prepared as a single cell suspension, and red blood cells were lysed for 1 minute using 1x RBC lysis buffer (Biolegend #420302). Cells were plated in M-CSF conditioned media: DMEM (Gibco) with 10% FBS, 1% PenStrep (Sigma Aldrich), 1% Sodium Pyruvate (Millipore), 1% L-glutamine (Sigma Aldrich), and 10% CMG 14-12 cell supernatant. Cells were maintained in a 37°C incubator with 5% CO_2_. Media was replenished on day 4. On day 7, cells were lifted from the plate using Trypsin-EDTA (Gibco) and used for *in vitro* assays. Macrophage identity was confirmed via flow cytometry based on high surface expression of F4/80 and CD11b.

### LAM and ExLAM Generation

For all *in vitro* assays, 200,000 cells were plated in each well of a 24-well plate and allowed to adhere at 37°C. To generate LAMs, BMDMs were treated overnight with 20 µg/mL soluble cholesterol (Sigma Aldrich #C4951). For ExLAMs, following lipid loading, wells were replaced with clean media and incubated overnight to induce cholesterol unloading. The next day, media was replaced again followed by an additional overnight incubation, for a total of 48 hours of lipid efflux. Lipid loading and unloading was confirmed via flow cytometry and fluorescent imaging with BODIPY-FL. Image quantification was performed in QuPath.

### Efferocytosis Assay

For efferocytosis assays, apoptosis of splenocytes was induced by irradiation at 33.3 Grays followed by 1.5 hours of rest in a 37°C incubator. Apoptosis was confirmed by flow cytometry using Annexin-V and propidium iodide (PI) staining. Apoptotic cells (ACs) were then labeled with CellTrace Violet (CTV, Life Technologies #C34557) and co-cultured with BMDMs (pre-treated with cholesterol, if applicable) in a 5:1 ratio for 1 hour at 37°C. After co-culture, cells were washed and stained for macrophage markers F4/80 and CD11b. Engulfment was assessed by flow cytometry where CTV+ BMDMs were quantified.

### Continual Efferocytosis Assay

As above, splenocytes were irradiated and apoptosis confirmed by flow cytometry. ACs were then split and labeled individually with CellTrace Violet (CTV) and CellTracker Green (CTG, Life Technologies #C7025). BMDMs were first incubated with CTV-labeled ACs (5 ACs:1 BMDM) for 45 minutes at 37°C. Cells were washed with 1X PBS and allowed to rest for 2 hours at 37°C. BMDMs were then incubated with CTG-labeled ACs for an additional 45 minutes at 37°C. Cells were washed with 1X PBS and then lifted with Trypsin-EDTA (Gibco) for flow cytometry.

### Competitive Efferocytosis Assay

WT and TREM2^CR^ BMDMs were labeled with CellTrace Violet (CTV) and CellTracker Green (CTG), respectively, and co-cultured in a 1:1 ratio in a 24-well plate (200k BMDMs total per well). As above, splenocytes were irradiated and apoptosis confirmed by flow cytometry. ACs were then labeled with CellTracker Orange (CTO, Life Technologies #C34551) and co-cultured with BMDMs (pre-treated with cholesterol, if applicable) in a 5:1 ratio for 1 hour at 37°C. Cells were then washed and engulfment was assessed by flow cytometry. WT and TREM2^CR^ cells were identified by CTV and CTG, respectively, and the percentage of CTO+ engulfed cells for each BMDM population was quantified.

### Lactate Dehydrogenase (LDH) Assay

Control, LAM, and ExLAM BMDMs were generated as described. Manufacturer’s instructions for the Cytotoxicity Detection Kit (Roche #11644793001) were followed. Briefly, 100 µl of culture supernatant was mixed with 100 µl freshly prepared reaction mixture. A media only control was also included. Samples were placed on a gentle shaker for 30 minutes at room temperature. Following incubation, absorbance at 490nm was measured by spectrophotometer.

### BrdU Incorporation

One day prior to analysis, BrdU was incubated with control, LAM, and ExLAM BMDMs overnight (eBioscience # 8812-6600-42). For the ExLAM condition, BrdU was added on the second day of cholesterol efflux only; all conditions were incubated with BrdU for an equal amount of time one day prior to analysis. Cells were then stained for surface markers, fixed and permeabilized using freshly prepared BrdU staining buffer (per manufacturer’s instructions), and DNase was applied for 1 hour at 37°C. Cells were then stained with anti-BrdU PE antibody at a 1:20 dilution for 30 minutes and analyzed by flow cytometry.

### Isolation of Peritoneal Macrophages

Peritoneal lavage was performed on euthanized mice using cold PBS. Peritoneal macrophages (PMACs) were negatively selected for by magnetic bead isolation using the following antibodies: biotin B220 (clone RA3-6B2, Biolegend #103203), biotin Ly6G (clone 1A8, Biolegend #127603), and biotin TCRβ (clone H57-597, Biolegend #109203). Isolated PMACs were adhered to chamber slides for 30 minutes, followed by fixation. Lipid levels were assessed via fluorescent imaging with LipidTox Deep Red (Invitrogen #H34477). Image quantification was performed in QuPath.

### Flow Cytometry

Single cell suspensions were washed in FACS buffer (HBSS with 2% FBS and 2% EDTA). Cell pellets were resuspended and stained in antibody cocktail for 45 minutes at 4°C. Samples were run on the BD LSRFortessa or the BD LSRFortessa X-20. All machines are maintained and supported by the UMN Flow Cytometry core facility. Data was analyzed in FlowJo software (TreeStar). The following antibodies were used: CD45 BV785 (clone 30-F11, Biolegend #103149); CD11b BV605 (clone M1/70, Biolegend #101257), Ly6G AF488 (clone 1A8, Biolegend #127626), TCRβ APC (clone H57-597, Biolegend #109212), CD19 PE (clone 6D5, Biolegend #115508), Ly6C AF700 (clone HK1.4, Biolegend #128024), Viability Stain Ghost Dye Violet 510 (Tonbo Biosciences #13-0870-T100), CD115 PE-Cy7 (clone AFS98, Biolegend #135524), TREM2 APC (R&D Systems #FAB17291A), F4/80 FITC (clone BM8, Biolegend #123108).

### Bulk RNA Sequencing of BMDMs

LAMs and ExLAMs were generated from differentiated BMDMs, as described above. Cells were lysed and stored in Trizol at -80°C until RNA isolation using the RNeasy Micro Kit (Qiagen #74004). RNA was quantified on a NanoDrop and samples with <10 ng total RNA were discarded. Sequencing was performed by the University of Minnesota Genomics Core (UMGC) using the AVITI24 platform and the Watchmaker stranded mRNA library prep kit. Raw data were processed using the CHURP pipeline developed by the Minnesota Supercomputing Institute, which implemented and integrated Trimmomatic, HISAT2, SAMTools and featureCounts. Data were aligned to the *Mus musculus* GRCm38 (Ensembl release 102) mouse reference genome. The output gene expression matrix was transferred to R for further analysis. Differential gene expression analysis was performed using the DESeq2 package and pathway analysis was performed using the fgsea package. Pathways utilized were from the Gene Ontology Biological Pathways (GOBP) library acquired using the msigdbr package.

### Statistics and Rigor

All statistics were performed in GraphPad Prism software or R. Comparison between two groups used a student’s t-test while groups of three or more used a one-way ANOVA. Error bars on graphs represent standard error of the mean (S.E.M). P values were considered statistically significant below 0.05. In graphs, * p < 0.05, ** p < 0.01, *** p < 0.001, and **** p < 0.0001. Each experiment was repeated at least two independent times, unless otherwise specified.

## ACKNOWLEDGEMENTS

We would like to thank Bruce Lamb (Indiana University) for sharing the *Trem2^flox^* mouse strain and Marco Colonna (Washington University in St. Louis, currently Mass General Research Institute) for sharing the *Trem2^-/-^* and TREM2-CR strains. We would like to thank the UMN Flow Cytometry core facility, UMN Genomics core facility, and the Minnesota Supercomputing Institute for assistance acquiring the data herein. Finally, we would like to thank Alector Therapeutics for providing the AL002a Trem2 agonist antibody that was utilized in these studies. Research was funded by grant support from the National Institutes of Health (NIH) R01 HL144843 (JWW), NIH R01 AI165553 (JWW), MN Partnership for Biotechnology award (JWW), F31 HL182131 (HH), and T32 HL166142 (MDC). HH was also supported by an American Heart Association predoctoral fellowship (AHA 25PRE1361476). GMS was supported by AHA 26POST1543363.

## DECLARATION

The authors declare no conflicts of interest related to this work.

## CONTRIBUTIONS

JWW, MC, TJB, and HH conceived and designed the research project. HH, GMS, SL, MTP, MDC, AEK, PRS, YZ, and OM performed experiments. HH and JWW wrote the manuscript and prepared figures. All authors assisted with data interpretation and manuscript review.

## REFERENCES

1. Sarraju, A. & Nissen, S. E. Atherosclerotic plaque stabilization and regression: a review of clinical evidence. Nat Rev Cardiol 21, 487–497 (2024).

2. Potteaux, S. et al. Suppressed monocyte recruitment drives macrophage removal from atherosclerotic plaques of Apoe-/-mice during disease regression. Journal of Clinical Investigation 10.1172/JCI43802 (2011) doi:10.1172/JCI43802.

3. Rahman, K. et al. Inflammatory Ly6Chi monocytes and their conversion to M2 macrophages drive atherosclerosis regression. Journal of Clinical Investigation 127, 2904–2915 (2017).

4. Nicholls, S. J. et al. Effect of Evolocumab on Progression of Coronary Disease in Statin-Treated Patients: The GLAGOV Randomized Clinical Trial. JAMA 316, 2373 (2016).

5. Pandit, R., Hillman, H., Williams, J. W. & Yurdagul, A. More Than a Cleanup Crew: The Expanding Biology of Efferocytosis. ATVB 46, (2026).

6. Mehrotra, P. & Ravichandran, K. S. Drugging the efferocytosis process: concepts and opportunities. Nat Rev Drug Discov 21, 601–620 (2022).

7. Jaitin, D. A. et al. Lipid-Associated Macrophages Control Metabolic Homeostasis in a Trem2-Dependent Manner. Cell 178, 686–698.e14 (2019).

8. Xu, R. et al. Lipid-associated macrophages between aggravation and alleviation of metabolic diseases. Trends in Endocrinology & Metabolism 35, 981–995 (2024).

9. De Ponti, F. F. et al. Spatially restricted and ontogenically distinct hepatic macrophages are required for tissue repair. Immunity 58, 362–380.e10 (2025).

10. Molgora, M. et al. TREM2 Modulation Remodels the Tumor Myeloid Landscape Enhancing Anti-PD-1 Immunotherapy. Cell 182, 886–900.e17 (2020).

11. Patterson, M. T. et al. Trem2 Agonist Reprograms Foamy Macrophages to Promote Atherosclerotic Plaque Stability—Brief Report. ATVB 44, 1646–1657 (2024).

12. Colonna, M. The biology of TREM receptors. Nat Rev Immunol 23, 580–594 (2023).

13. Patterson, M. T. et al. Trem2 promotes foamy macrophage lipid uptake and survival in atherosclerosis. Nat Cardiovasc Res 2, 1015–1031 (2023).

14. Piollet, M. et al. TREM2 protects from atherosclerosis by limiting necrotic core formation. Nat Cardiovasc Res 3, 269–282 (2024).

15. Williams, J. W. et al. Limited macrophage positional dynamics in progressing or regressing murine atherosclerotic plaques—brief report. Arteriosclerosis, thrombosis, and vascular biology 38, 1702–1710 (2018).

16. Komukai, K. et al. Effect of Atorvastatin Therapy on Fibrous Cap Thickness in Coronary Atherosclerotic Plaque as Assessed by Optical Coherence Tomography. Journal of the American College of Cardiology 64, 2207–2217 (2014).

17. Gerlach, B. D. et al. Efferocytosis induces macrophage proliferation to help resolve tissue injury. Cell Metabolism 33, 2445–2463.e8 (2021).

18. Kim, K. W., Ivanov, S. & Williams, J. W. Monocyte Recruitment, Specification, and Function in Atherosclerosis. Cells 10, (2020).

19. Xu, Y. et al. Identification of conserved and tissue-restricted transcriptional profiles for lipid associated macrophages. Commun Biol 8, 953 (2025).

20. Lin, J.-D. et al. Single-cell analysis of fate-mapped macrophages reveals heterogeneity, including stem-like properties, during atherosclerosis progression and regression. JCI Insight 4, e124574, 124574 (2019).

21. Schlepckow, K. et al. An Alzheimer-associated TREM2 variant occurs at the ADAM cleavage site and affects shedding and phagocytic function. EMBO Mol Med 9, 1356–1365 (2017).

22. Ewers, M. et al. Increased soluble TREM2 in cerebrospinal fluid is associated with reduced cognitive and clinical decline in Alzheimer’s disease. Sci. Transl. Med. 11, eaav6221 (2019).

23. Kim, S.-M. et al. TREM2 promotes Aβ phagocytosis by upregulating C/EBPα-dependent CD36 expression in microglia. Sci Rep 7, 11118 (2017).

24. Chen, J. et al. GLP-1 receptor agonist as a modulator of innate immunity. Front. Immunol. 13, 997578 (2022).

25. Yusta, B. et al. GLP-1 receptor activation improves β cell function and survival following induction of endoplasmic reticulum stress. Cell Metabolism 4, 391–406 (2006).

26. Caslin, H. L., Cottam, M. A., Piñon, J. M., Boney, L. Y. & Hasty, A. H. Weight cycling induces innate immune memory in adipose tissue macrophages. Front. Immunol. 13, 984859 (2023).

27. Wang, H., He, W., Yang, G., Zhu, L. & Liu, X. The Impact of Weight Cycling on Health and Obesity. Metabolites 14, 344 (2024).

28. Delahanty, L. M. et al. Effects of Weight Loss, Weight Cycling, and Weight Loss Maintenance on Diabetes Incidence and Change in Cardiometabolic Traits in the Diabetes Prevention Program. Diabetes Care 37, 2738–2745 (2014).

29. Takaoka, M. et al. Early intermittent hyperlipidaemia alters tissue macrophages to fuel atherosclerosis. Nature 634, 457–465 (2024).

30. Lavillegrand, J.-R. et al. Alternating high-fat diet enhances atherosclerosis by neutrophil reprogramming. Nature 634, 447–456 (2024).

31. Zhang, Q. et al. Landscape and Dynamics of Single Immune Cells in Hepatocellular Carcinoma. Cell 179, 829–845.e20 (2019).

32. Katzenelenbogen, Y. et al. Coupled scRNA-Seq and Intracellular Protein Activity Reveal an Immunosuppressive Role of TREM2 in Cancer. Cell 182, 872–885.e19 (2020).

33. Timperi, E. et al. Lipid-Associated Macrophages Are Induced by Cancer-Associated Fibroblasts and Mediate Immune Suppression in Breast Cancer. Cancer Research 82, 3291–3306 (2022).

34. Masetti, M. et al. Lipid-loaded tumor-associated macrophages sustain tumor growth and invasiveness in prostate cancer. Journal of Experimental Medicine 219, e20210564 (2022).

35. Yang, P. et al. CD36-mediated metabolic crosstalk between tumor cells and macrophages affects liver metastasis. Nat Commun 13, 5782 (2022).

36. Croxford, A. L. et al. The Cytokine GM-CSF Drives the Inflammatory Signature of CCR2+ Monocytes and Licenses Autoimmunity. Immunity 43, 502–514 (2015).

