## Supplemental Figures S1-S8 for "TREM2 encodes plaque macrophage plasticity to mediate atherosclerosis regression"

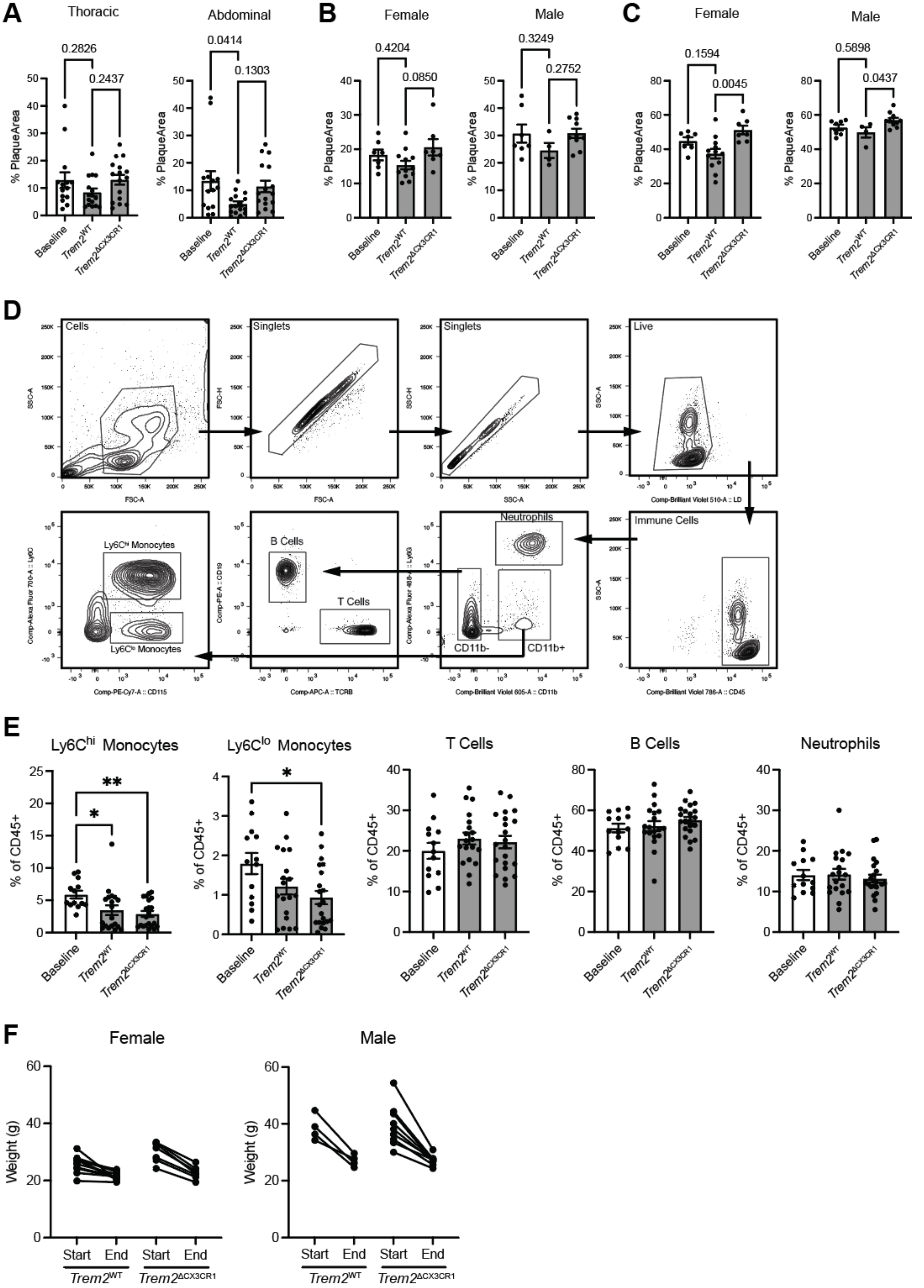


**Figure S1. *Trem2* deletion in CX3CR1-expressing cells has no impact on circulating immune cells or body weight.** A) Quantification of plaque area in the thoracic and abdominal aorta regions in baseline and regression mice. B) Total plaque area and C) arch plaque area in males and females included in the regression study. D) Flow cytometry gating scheme to identify major immune cell populations in blood and E) quantification of blood immune cells in mice one day prior to euthanasia. F) Body weight (grams) of *Trem2*$^{WT}$ and *Trem2*$^{\Delta CX3CR1}$ mice at the start and end of the regression period.

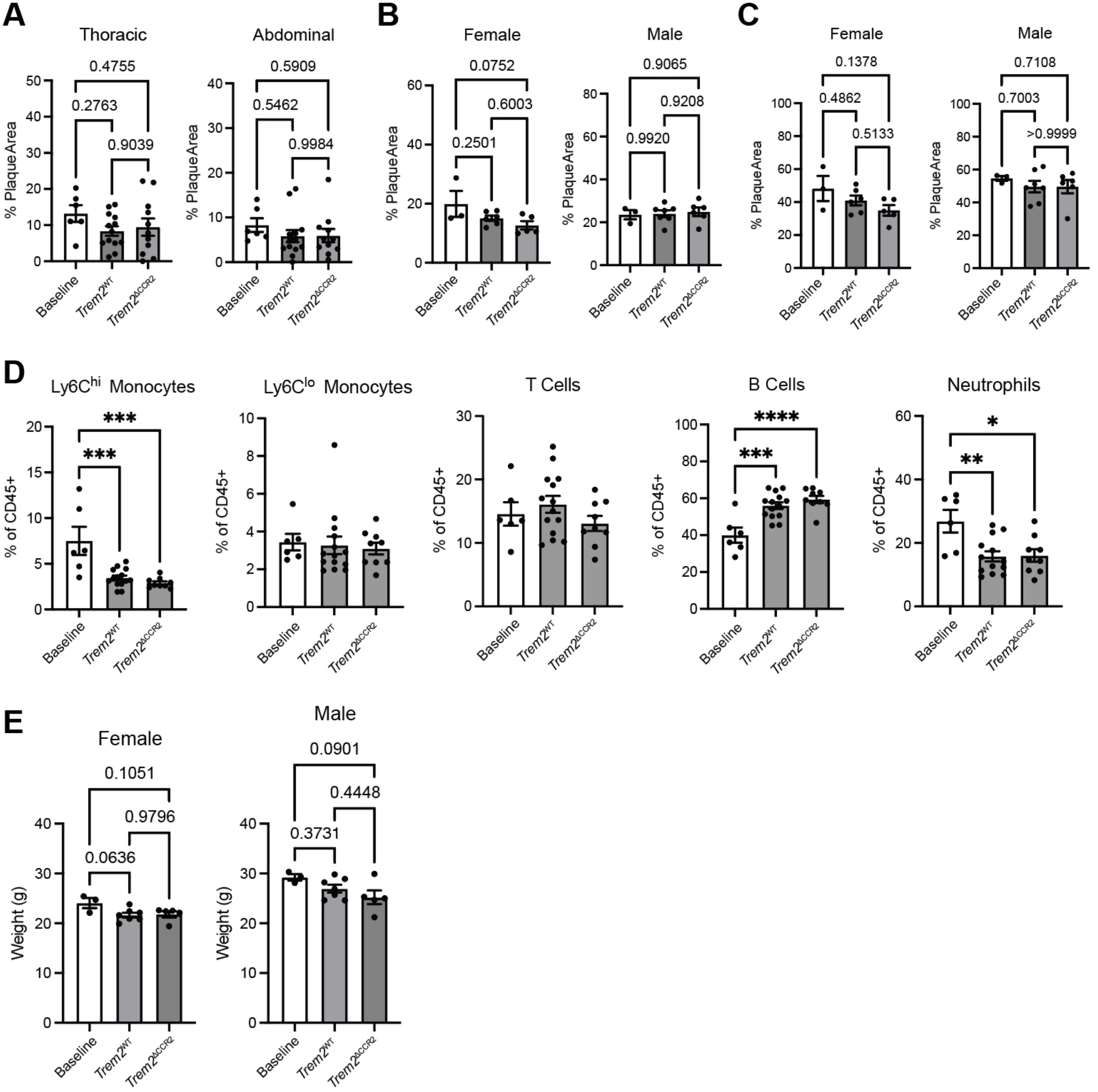


**Figure S2. *Trem2* deletion in CCR2-expressing cells has no impact on circulating immune cells or body weight.** A) Quantification of plaque area in the thoracic and abdominal aorta regions in regression mice. B) Total plaque area and C) arch plaque area in males and females included in the regression study. D) Quantification of blood immune cells in mice one day prior to euthanasia. The same gating scheme as Figure S6D was used. F) Body weight (grams) of Baseline, *Trem2*$^{WT}$, and *Trem2*$^{\Delta CCR2}$ mice at the experimental end point.

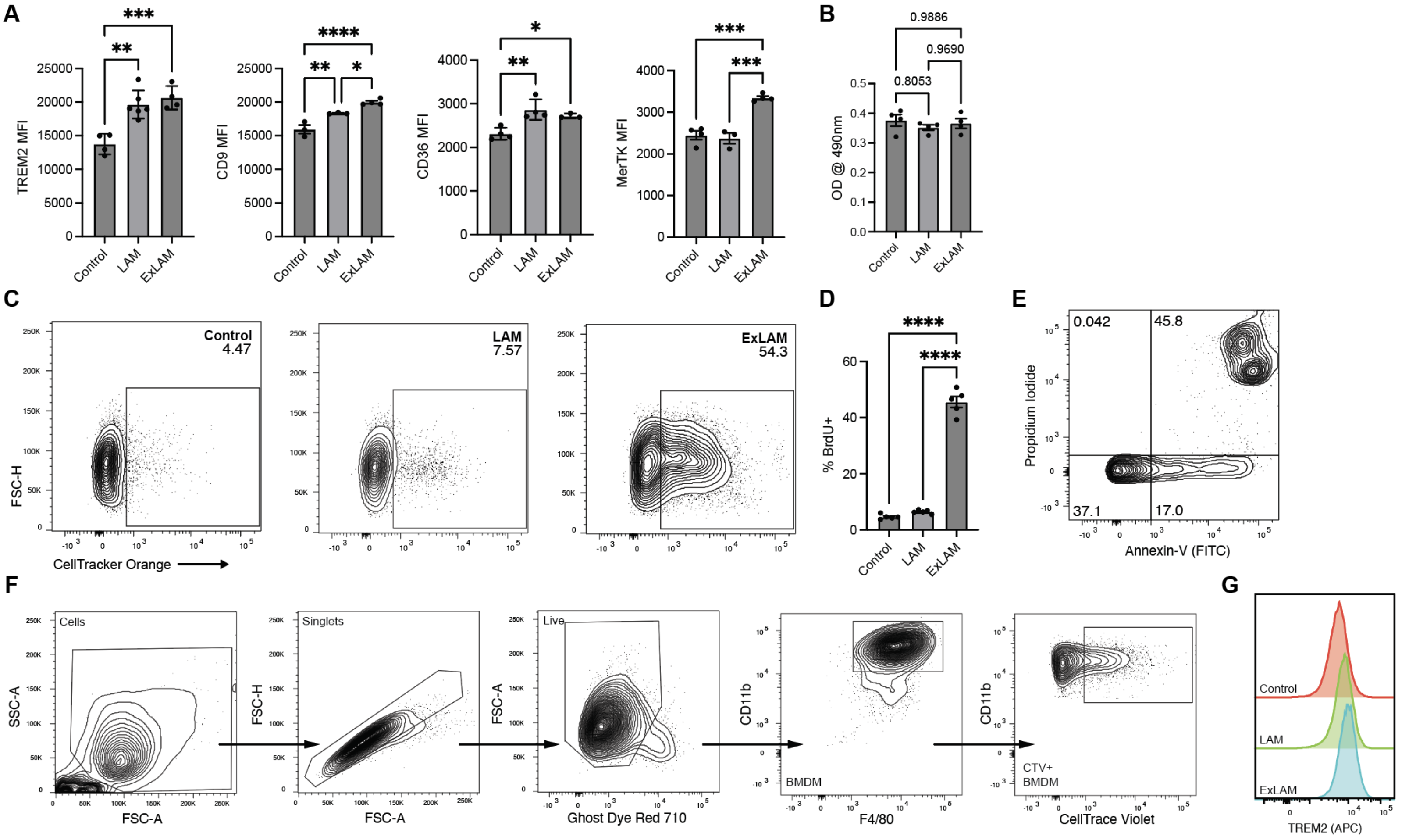


**Figure S3. ExLAMs maintain surface expression of LAM markers and have enhanced efferocytic ability.** A) Mean fluorescence intensity (MFI) of LAM signature proteins in control, LAMs, and ExLAMs, as measured by flow cytometry. B) Cell death measured by LDH release assay. C) Representative contour flow cytometry plots and D) quantification of BMDM proliferation as measured by BrdU incorporation. E) Annexin-V and propidium iodide (PI) staining for apoptotic cells by flow cytometry. Apoptosis of irradiated splenocytes was confirmed for efferocytosis assays. F) Flow cytometry gating scheme for efferocytosis experiments. G) Histograms of TREM2 mean fluorescent intensity (quantification in Figure 6F).

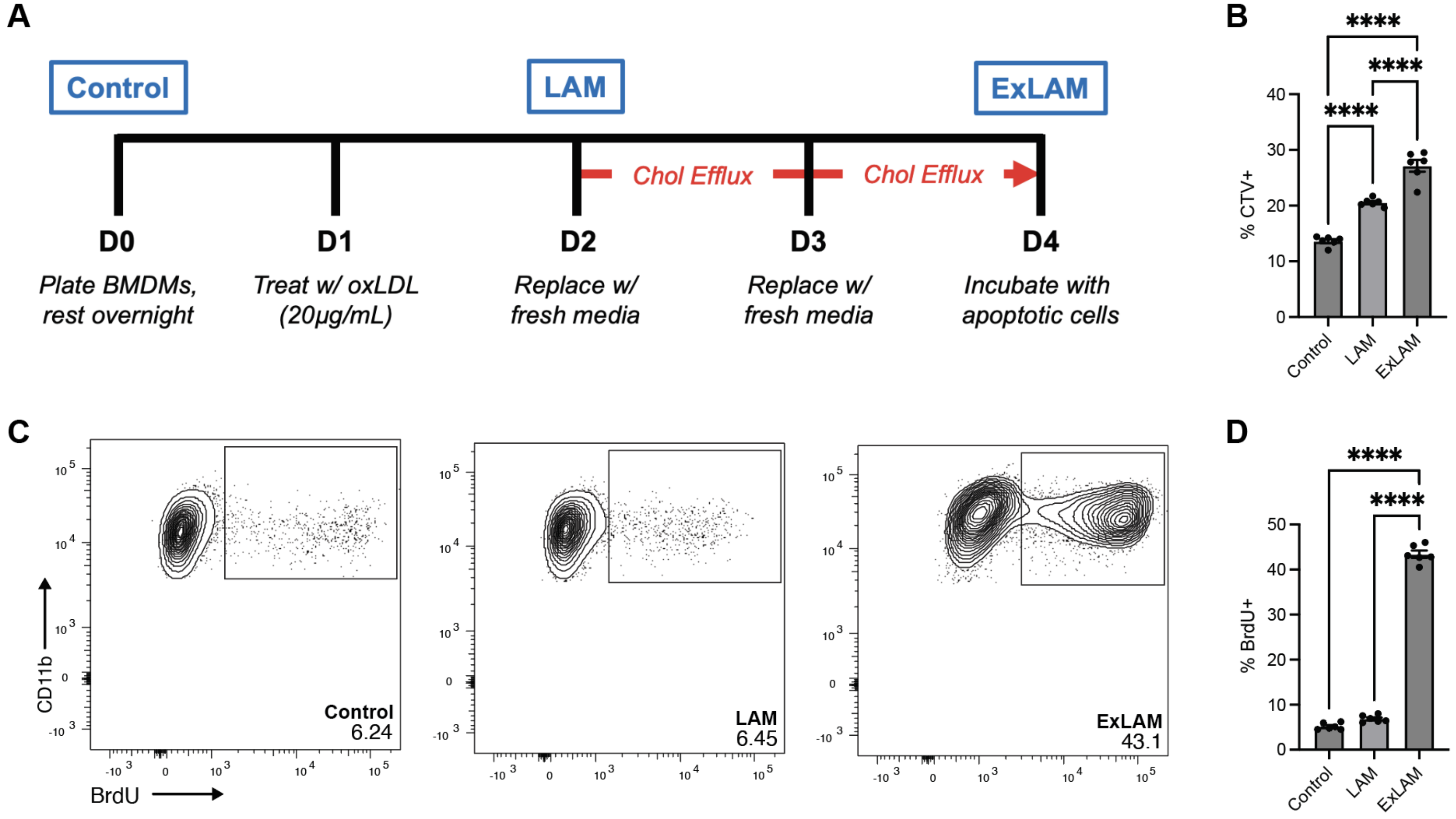


**Figure S4. Enhanced efferocytosis and proliferation are recapitulated in oxLDL-derived ExLAMs.** A) Experimental setup for ExLAM cell generation with oxidized low-density lipoprotein (oxLDL). Briefly, ExLAMs were loaded with 20µg/mL oxLDL overnight to become LAMs and then allowed to efflux cholesterol over a two-day period before being used in assays. LAMs (20µg/mL oxLDL overnight) and Control (no cholesterol) cells are used as controls. B) Quantification of oxLDL BMDMs (CD11b+ F4/80+) that have engulfed a CellTrace Violet (CTV)-labeled apoptotic splenocyte. C) Representative contour flow cytometry plots and D) quantification of BMDM proliferation as measured by BrdU incorporation.

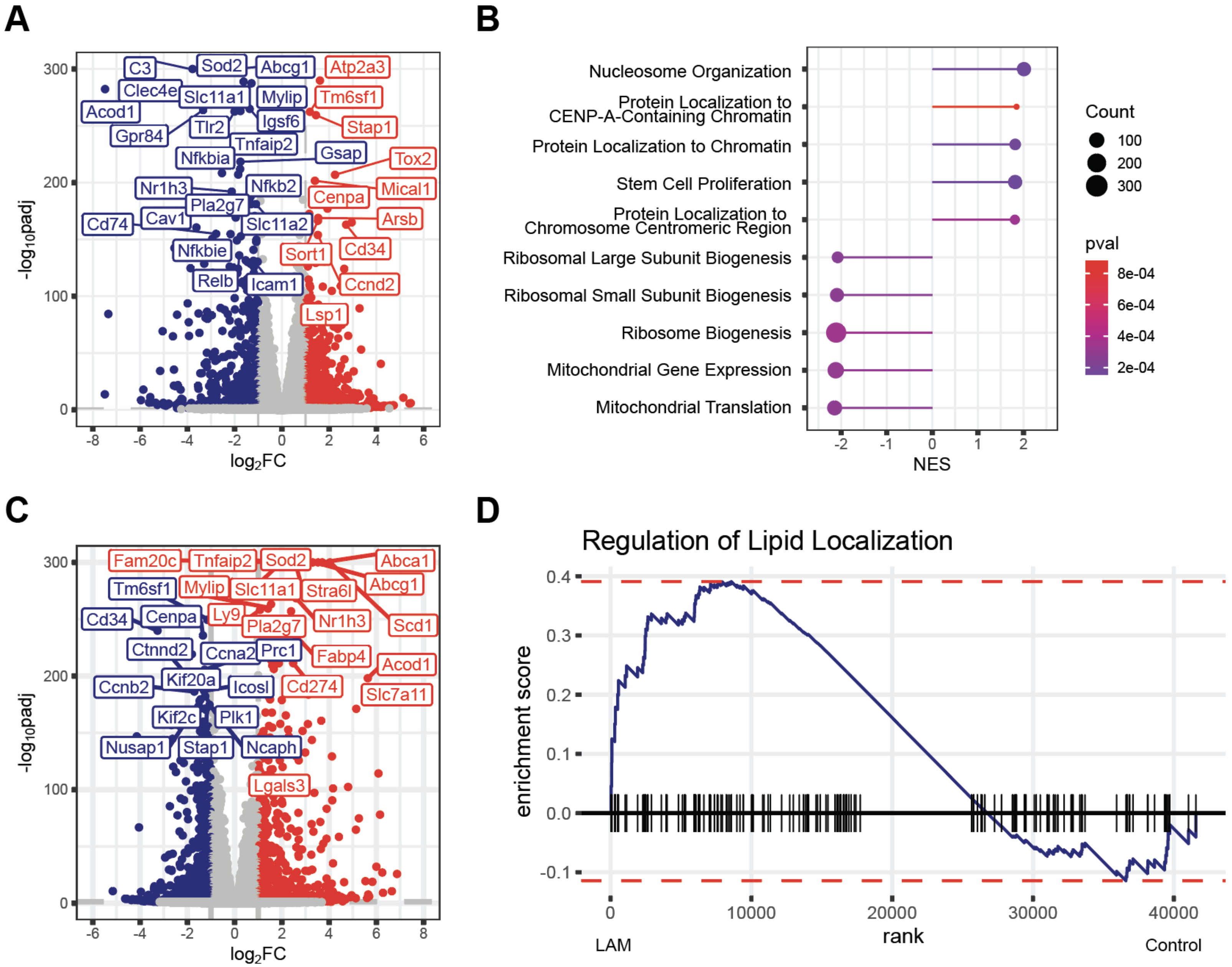


**Figure S5. Bulk RNA sequencing of ExLAMs reveals upregulation of chromatin genes compared to LAMs.** A) Volcano plot of top DEGs between ExLAMs vs LAMs. padj <0.05 and abs($Log_2FC$ > 1) was considered significant. Genes in red are significantly upregulated in ExLAMs and genes in blue are significantly downregulated. B) Pathway analysis using DEGs from (A). Gene Ontology Biological Processes (GOBP) pathway library was utilized. C) Volcano plot of top DEGs between LAMs vs control BMDMs. padj <0.05 and abs($Log_2FC$) > 1 was considered significant. Genes in red are significantly upregulated in ExLAMs and genes in blue are significantly downregulated. D) Gene set enrichment analysis (GSEA) of Regulation of Lipid Localization pathway (GOBP) in LAMs vs control.

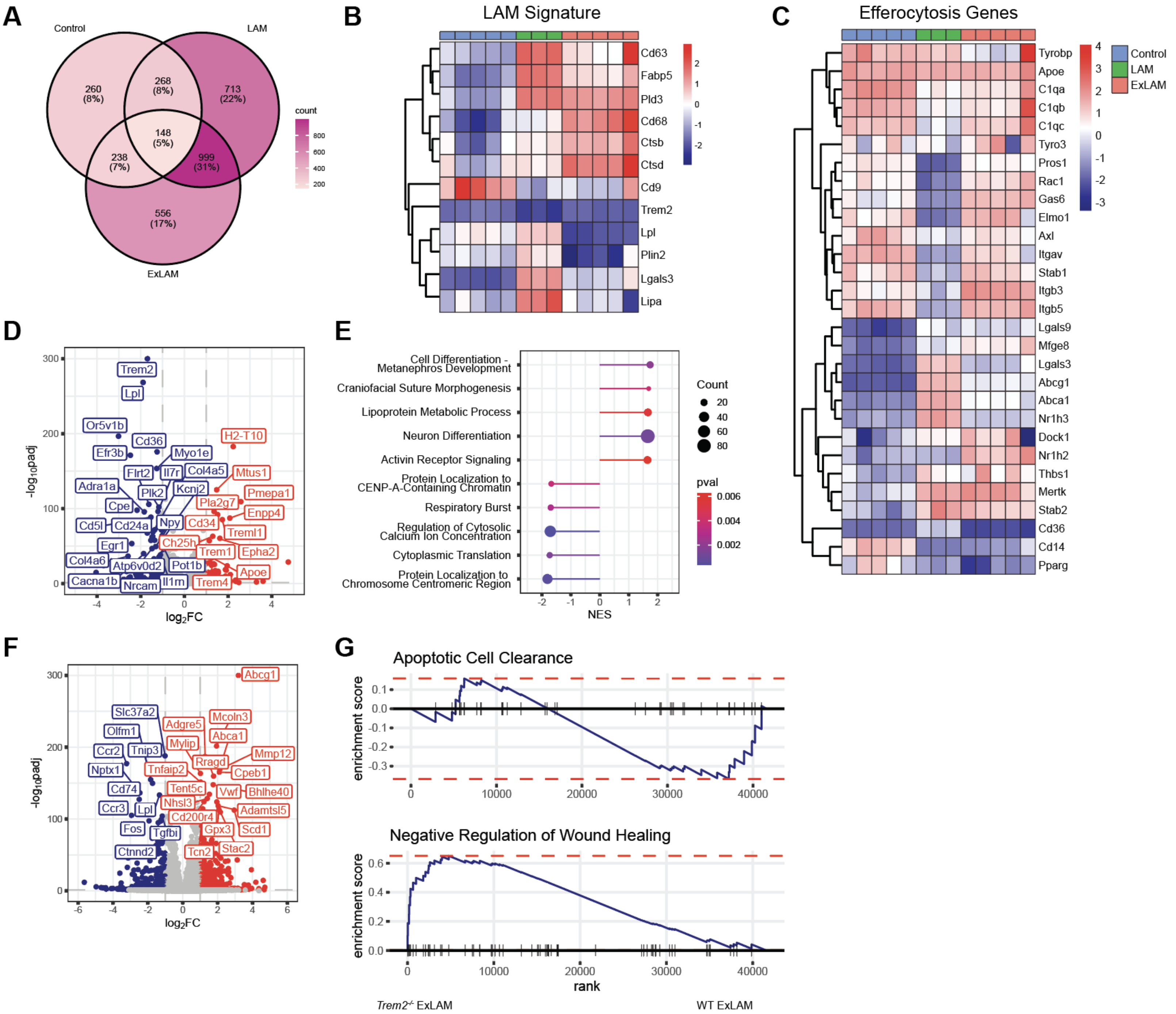


**Figure S6. *Trem2*-deficient ExLAMs fail to upregulate efferocytosis and wound healing genes.** A) Venn diagram showing the number of differentially-expressed genes (DEGs) in *Trem2*$^{-/-}$ control, LAMs, and ExLAMs. B) Heat map of relative expression of lipid-associated macrophage genes across the three treatment groups. Data shown are row-wise Z scores. C) Heat map of relative expression of efferocytosis-associated genes across the three treatment groups. Data shown are row-wise Z scores. D) Volcano plot of differentially expressed genes between *Trem2*$^{-/-}$ ExLAMs and WT ExLAMs. padj <0.05 and abs($Log_2$FC) > 1 was considered significant. Genes in red are significantly upregulated in *Trem2*$^{-/-}$ ExLAMs cells and genes in blue are significantly downregulated. E) Pathway analysis using DEGs from (D). Gene Ontology Biological Processes (GOBP) pathway library was utilized. F) Volcano plot of differentially expressed genes between *Trem2*$^{-/-}$ ExLAMs and Trem2$^{-/-}$ controls. padj <0.05 and abs($Log_2$FC) > 1 was considered significant. Genes in red are significantly upregulated in *Trem2*$^{-/-}$ ExLAMs cells and genes in blue are significantly downregulated. G) Gene set enrichment analysis (GSEA) of pathways in *Trem2*$^{-/-}$ ExLAMs vs WT ExLAMs. A peak to the left of the graph indicates upregulation of genes in the given pathway in *Trem2*$^{-/-}$ ExLAMs, while a peak to the right indicates upregulation in WT ExLAMS.

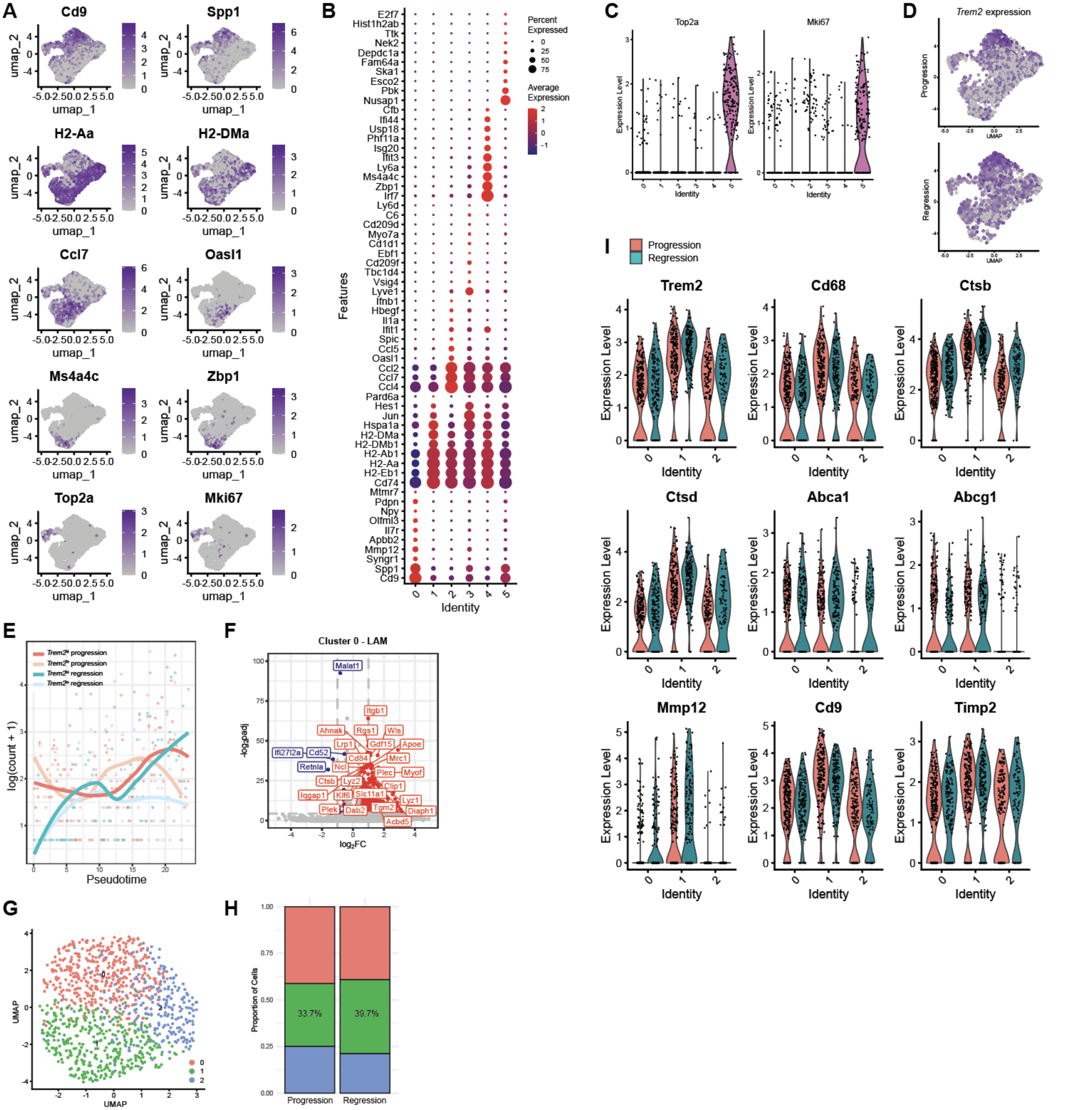

**Figure S7. Murine aortic plaques contain an ExLAM-like macrophage subcluster that upregulate lysosomal and lipid processing genes.** A) Individual gene FeaturePlots for cluster identification. B) DotPlot of top 10 cluster-defining genes for each cluster in the UMAP. C) *Top2a* and *Mki67* expression in each cluster of the UMAP. D) *Trem2* expression projected onto the UMAP. E) Pseudotime (Monocle3) of all aortic cells split by *Trem2* expression (high vs low) and experimental group (progression vs regression). Root node was manually selected to be the monocyte cluster 4 from UMAP in Figure 6G. *Trem2* expression modulates macrophage fate prediction along differentiation trajectory from monocyte origin. F) Volcano plot of top DEGs between Cluster 0 in Regression vs Cluster 0 in Progression. padj <0.05 and abs($Log_2FC$) > 1 was considered significant. Genes in red are significantly upregulated in regression and genes in blue are significantly downregulated. G) LAM cluster 0 from Figure 6G was reclustered to identify three LAM subclusters. H) Cluster proportions from (G) between the two experimental groups. I) Select genes from ExLAM *in vitro* bulk RNA sequencing used to identify the ExLAM population in the single cell dataset.

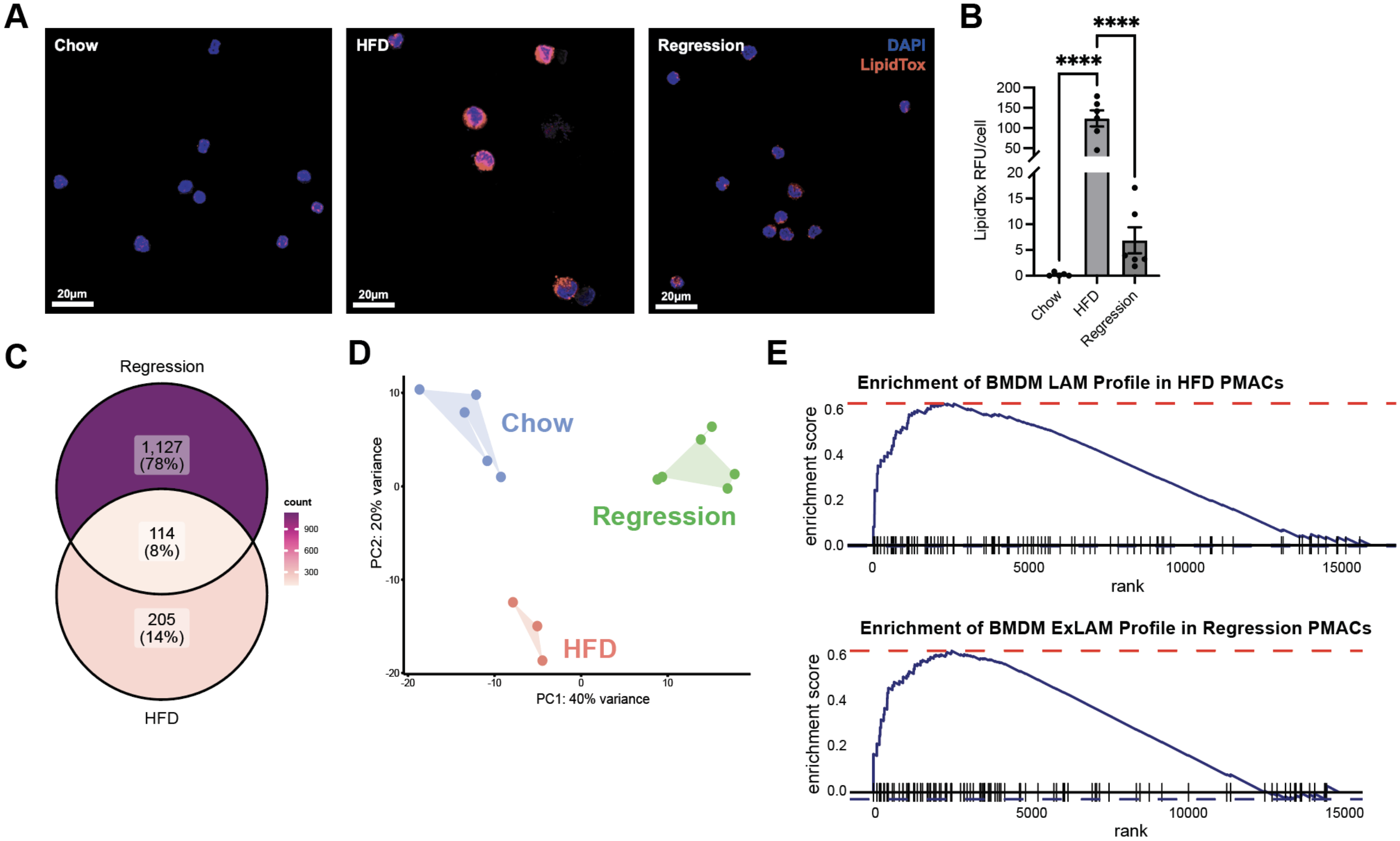


**Figure S8. Peritoneal macrophages in a mouse model of atherosclerosis regression transcriptionally resemble *in vitro*-generated ExLAMs.** A) Immunofluorescent imaging of isolated peritoneal macrophages (PMACs) stained for nuclei (DAPI) and lipid content (LipidTox). B) Quantification of (A). C) Venn diagram showing the number of differentially-expressed genes (DEGs) in PMACs from high fat diet (HFD)-fed and regression mice, as compared to chow-fed mice. D) Principal component analysis (PCA) plot of bulk RNA sequencing data of mouse PMACs. E) Gene set enrichment analysis (GSEA) comparing top 100 DEGs in BMDM LAMs vs PMACs from HFD-fed mice (top) and BMDM ExLAMs vs PMACs from regression mice (bottom). A peak to the left of the graph indicates high similarity between BMDM and PMAC top 100 genes, while a peak to the right of the graph indicates low similarity in gene expression.
